# IsoDGR-Induced Endothelial Cytoskeletal Disruption Drives Age-Related Blood-Brain Barrier Breakdown

**DOI:** 10.1101/2025.07.08.663800

**Authors:** Ridhi Patel, Pazhanichamy Kalailingam, SoFong Cam Ngan, Ranjith Iyappan, Jones Nwaeze, Bianca D’Amore, Maryam Katoueezadeh, Yazhini Thinakaran, Mahdi Hejazi Laein, Brent E. Faught, Rebecca E. K. Macpherson, Evangelia Litsa Tsiani, Panagiota Klentrou, Dominique de Kleijn, Raj N. Kalaria, Neil E. McCarthy, Siu Kwan Sze

**Author notes:** **Corresponding authors** Siu Kwan SZE, PhD, Department of Health Sciences, Faculty of Applied Health Sciences, Brock University, St. Catharines, Ontario L2S 3A1, Canada. These authors contributed equally to this work (Ridhi Patel, Pazhanichamy Kalailingam).

## Abstract

Brain aging is characterized by progressive breakdown of the blood-brain barrier (BBB), which correlates with neuroinflammation and cognitive decline. Emerging evidence implicates degenerative modifications of the vascular proteins as a key driver of BBB dysfunction. In particular, spontaneous deamidation of Asp-Gly-Arg (NGR) motifs generates isoAsp-Gly-Arg (isoDGR) sequences that structurally mimic canonical Arg-Gly-Asp (RGD) integrin-binding ligands. Here, we show that age-associated accumulation of isoDGR in the brain cortex induces endothelial cytoskeletal collapse and tight junction disorganization, leading to BBB breakdown. Using mice lacking the L-isoaspartyl repair enzyme PCMT1 (which accelerates isoDGR accumulation) and wild type aged mice, we found markedly elevated isoDGR in brain tissues accompanied by focal microhemorrhages and increased BBB permeability. Recent whole-genome sequencing suggests that a common PCMT1 variant is linked to neurodegenerative disease risk, indicating potential clinical relevance in vascular aging. Remarkably, systemic treatment with an isoDGR-neutralizing antibody largely prevented capillary breaches and leakage, and even restored barrier integrity in aged wild-type mice. To uncover the molecular mechanism, we exposed brain endothelial cells to synthetic isoDGR-peptides, which recapitulated these effects. Unbiased RNA-sequencing reinforced these findings, revealing broad transcriptomic reprogramming of cytoskeletal, cell-cell junction, inflammatory, and stress-response pathways. Functional studies demonstrated that isoDGR triggered collapse of F-actin stress fibers, disrupted junctional ZO-1 and VE-cadherin, increased monolayer permeability to macromolecules, and impaired endothelial cell migration and proliferation. IsoDGR-treated endothelial cells exhibited increased oxidative stress, upregulation of ICAM-1/VCAM-1/CCL-2, and adopted a senescent phenotype. Our results suggest that isoDGR hijacks endothelial integrin signaling to destabilize the actin cytoskeleton and tight junctions, a process that breaches the BBB and subsequently activates inflammatory and senescence programs. In summary, we identify BBB disruption via isoDGR-induced cytoskeletal dysfunction as a central pathology of vascular aging, and demonstrate that targeting isoDGR damage preserves BBB integrity and attenuates neuroinflammation.

## INTRODUCTION

Progressive endothelial dysfunction and BBB breakdown are hallmarks of vascular aging and are strongly linked to neuroinflammation and cognitive decline^1–3^. While multiple factors contribute to age-related vascular deterioration, accumulating molecular damage in long-lived proteins has emerged as a critical mechanism^4–6^. Structural matrix components such as fibronectin, laminin, and collagens turn over slowly and accrue various degenerative protein modifications (DPMs) that alter their intrinsic biochemical properties over time^6–13^. One key age-associated DPM is asparagine deamidation, which can convert Asn-Gly-Arg (NGR) sequences into isoAsp-Gly-Arg (isoDGR) motifs^14–19^. IsoDGR is of particular interest because it closely mimics the Arg-Gly-Asp (RGD) motif that serves as a principal recognition motif for integrin receptors^20–22^.

Integrins are transmembrane adhesion receptors on endothelial cells that normally bind RGD sequences in ECM proteins to mediate cell attachment, migration, and intracellular signaling^23^. Importantly, integrins physically link the extracellular matrix to the actin cytoskeleton via focal adhesion complexes^24,25^. Thus, the formation of isoDGR damage motifs in aging ECM may aberrantly engage endothelial integrins and destabilize the cytoskeletal architecture. Indeed, our previous studies and others have shown that isoDGR-modified fibronectin can bind integrin receptors^26–32^ on endothelial and immune cells to trigger “outside-in” signaling, leading to activation of ERK/AP-1 inflammatory pathways, cytokine release (e.g. TNFα), and enhanced monocyte adhesion^16–19,33–35^. Moreover, chronic exposure to isoDGR has been shown to upregulate endothelial integrin β1 expression, potentially creating a feed-forward loop of increased cell-ECM adhesion^17^. Together, these findings suggest that isoDGR is a gain-of-function matrix lesion that converts the aged ECM into a pro-inflammatory and barrier-disrupting platform.

Protein L-isoaspartyl O-methyltransferase 1 (PCMT1) is an enzyme that repairs isoaspartate residues in proteins^36,37^, thereby limiting the accumulation of isoDGR damage. Accordingly, mice lacking PCMT1 (Pcmt1^−/-^) exhibit accelerated buildup of isoAsp and isoDGR in tissues, resulting in severe multiorgan pathology and premature death by ∼1-2 months^16–19,33,34,36,37^. In the vasculature, Pcmt1^−/-^ mice accumulate isoDGR-modified matrix proteins within blood vessels, at levels and distribution patterns comparable to those seen in older wild-type mice^17,18^ and in patients with cardiovascular disease^34^. Importantly, recent large-scale whole-genome sequencing in humans has revealed that individuals carrying a common 619 bp deletion in the PCMT1 gene have an 8.38-fold higher risk of developing neurodegenerative disease^38^, further underscoring the potential clinical significance of PCMT1 deficiency and isoDGR accumulation in age-related vascular pathology. However, the impact of age-related isoDGR accumulation on BBB integrity and brain health remains poorly understood. Given that BBB breakdown in normal aging is associated with neuroinflammation and cognitive impairment^1–3^, we hypothesized that isoDGR deposition in the aging neurovascular unit is a key driver of BBB dysfunction *in-vivo*.

In this study, we provide evidence that isoDGR accumulation triggers integrin-mediated signaling in brain endothelial cells, leading to cytoskeletal disorganization, loss of intercellular junctions, and consequent increases in vascular permeability and BBB disruption. The sustained engagement of these integrin and cytoskeletal pathways may provoke oxidative stress and pushes endothelial cells toward a premature senescent state. To test these concepts, we first examined isoDGR-linked BBB disruption and inflammation in *in-vivo* models including Pcmt1^−/-^ mice (which model accelerated isoDGR accumulation) and naturally aged wild-type mice, with or without treatment using an isoDGR-neutralizing antibody. We then used cultured brain microvascular endothelial cells treated with synthetic isoDGR-containing peptides to identify specific cellular and molecular changes induced by this motif *in-vitro*. An unbiased RNA-sequencing analysis was performed to capture the global transcriptomic effects of isoDGR on the endothelial cells. The isoDGR-induced changes were further validated by various functional assays. Our findings reveal a mechanistic pathway whereby age-related ECM damage (isoDGR) destabilizes the endothelial cytoskeleton and tight junctions, igniting chronic inflammatory and stress signaling that culminates in BBB breakdown and endothelial senescence. These results extend the paradigm of molecular aging damage to the neurovascular context and identify the isoDGR-integrin axis as a potential therapeutic target for preserving BBB function in the aging brain.

## RESULTS

### IsoDGR accumulation precipitates neurovascular injury and BBB leakage *in-vivo*

Brains of Pcmt1-knockout (Pcmt1^−/-^) mice, which lack the protein-repair enzyme PCMT1, contained dramatically higher levels of isoDGR-modified proteins in the cerebral cortex compared to age-matched wild-type controls (Fig. 1A,C). Immunofluorescence staining revealed dense isoDGR accumulation throughout both cortical and subcortical regions of Pcmt1^−/-^ brains, whereas wild-type mice exhibited only sparse isoDGR staining. This biochemical burden was associated with structural damage to the neurovascular unit. High-magnification imaging of capillaries (stained for the endothelial marker GLUT1) in Pcmt1^−/-^ brains showed focal breaches in the capillary wall, with extravasation of TER-119^+^ red blood cells into the surrounding parenchyma (Fig. 1B). Quantitative analysis confirmed a significant increase in both isoDGR signal intensity and the frequency of such microhemorrhages in cortex of Pcmt1^−/-^ versus wild-type brains (Fig. 1C,D), linking isoDGR accumulation to compromised vessel integrity.

**Figure 1.**
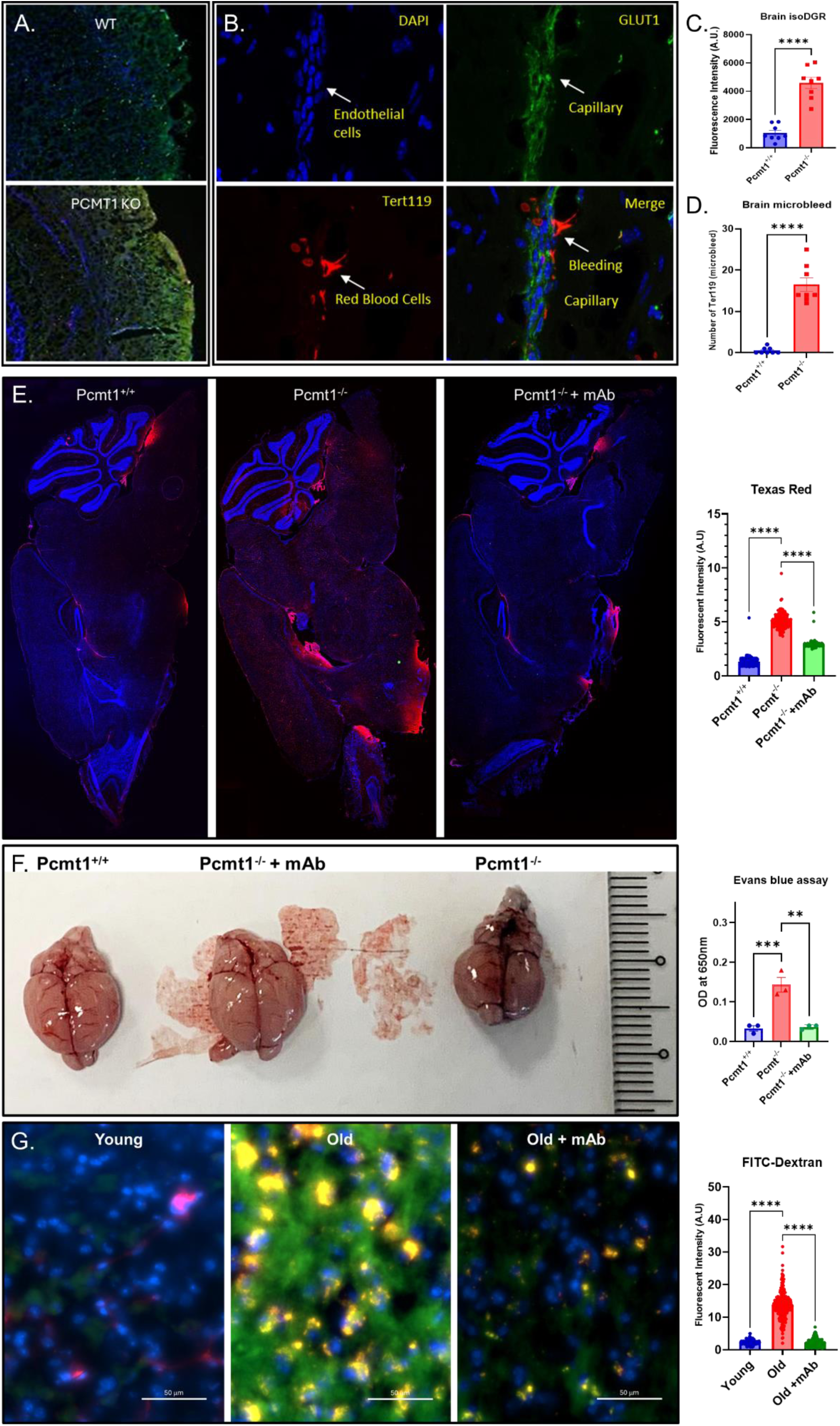
IsoDGR accumulation drives blood-brain barrier leakage in both Pcmt1-KO and wild type aged mice, and isoDGR-neutralizing antibody restores barrier integrity. (A) Representative cortical sections from 8-week-old wild-type (WT) and Pcmt1-knockout (KO) mice immunostained for isoDGR (green) with nuclei counterstained by DAPI (blue). The Pcmt1-KO brain shows diffuse isoDGR deposition throughout the parenchyma, whereas the age-matched WT brain exhibits only minimal isoDGR signal. (B) High-magnification images of KO brain microvessels stained for the endothelial marker GLUT1 (green) and red blood cells (TER-119, red). White arrows indicate focal capillary breaches with extravasated TER-119^+^ cells, illustrating microhemorrhages in the KO mouse (these lesions are absent in WT). (C) Quantification of isoDGR immunofluorescence intensity in cortical sections, confirming significantly higher isoDGR levels in Pcmt1-KO vs. WT brains. (D) Frequency of microhemorrhages (TER-119^+^ extravasations per mm²) in WT and KO brains. Pcmt1-KO mice show an increased incidence of microhemorrhages, consistent with compromised vessel integrity. (E) BBB tracer leakage in Pcmt1-KO vs. WT: Whole-brain sections 2 hours after intravenous injection of a high-molecular-weight tracer (70 kDa Texas Red-dextran). The KO brain displays widespread parenchymal red fluorescence, indicating extensive BBB leakage, whereas the WT control retains the tracer within the vasculature. Notably, KO mice pre-treated with isoDGR-neutralizing antibody (isoDGR-mAb, 1 mg/kg/week for 8 weeks before the tracer) show markedly reduced dextran extravasation, demonstrating that the BBB defect in KO is rescued by isoDGR neutralization. (F) Evans Blue extravasation assay: Representative brains collected 30 min after Evans Blue injection (which binds to albumin) from WT, KO, and isoDGR-mAb-treated KO mice. Brains from untreated KO mice are diffusely blue, reflecting substantial albumin leakage across the BBB, whereas WT brains remain largely unstained. IsoDGR-mAb treatment greatly diminishes Evans Blue leakage in KO mice, consistent with restored barrier function. Quantification (bar graph) of Evans Blue content in brain homogenates (absorbance at 650 nm) confirms a significant reduction in BBB permeability in KO mice receiving the antibody. (G) Age-associated BBB leakage: Twenty-six-month-old (aged) WT mice exhibit robust BBB leakiness compared to 3-month-old young adult WT mice. A small tracer (4 kDa FITC-dextran, green) was used in aged animals to detect subtle BBB disruptions. Fluorescent imaging reveals substantial parenchymal dextran accumulation in aged WT brains (indicative of BBB breakdown) relative to negligible leakage in young adults. Strikingly, isoDGR-mAb treatment in aged mice (same regimen as in KO) dramatically reduces FITC-dextran extravasation, restoring BBB integrity to near-youthful levels. Quantified fluorescence intensity in brain parenchyma highlights the significant protection conferred by isoDGR neutralization in aging. Fluorescence intensity was quantified using ImageJ software by measuring mean signal intensity in defined parenchymal regions. All quantitative data are presented as mean ± SEM. **p < 0.01, ****p < 0.0001 (unpaired t-test or one-way ANOVA with Tukey’s post hoc test).

Functional tracer assays corroborated these histological signs of BBB leakage. Following intravenous injection of fluorescent 70 kDa dextran, Pcmt1^−/-^ mice displayed widespread parenchymal fluorescence, indicating leakage of the tracer out of vessels, whereas wild-type animals retained the Texas-Red dye within the vascular lumen (Fig. 1E). Strikingly, systemic administration of an isoDGR-neutralizing monoclonal antibody (isoDGR-mAb, 1 mg/kg/week for 8 weeks before the tracer) markedly reduced dextran leakage in Pcmt1^−/-^ mice, demonstrating that the vascular permeability defect was isoDGR-dependent. A complementary macroscopic assay using Evans Blue (which binds albumin)^39^ yielded concordant results, Pcmt1^−/-^ brains were diffusely blue (albumin extravasation) whereas wild-type brains remained largely unstained, and isoDGR antibody treatment significantly attenuated Evans Blue leakage in the knockouts (Fig. 1F). Finally, to assess whether isoDGR contributes to BBB impairment during normal aging, we evaluated BBB integrity in naturally aged wild type mice. A small tracer (4 kDa FITC-dextran, green) was used in aged animals to detect subtle BBB disruptions^40,41^. Twenty-six-month-old wild-type mice showed robust tracer leakage compared to young adults, confirming an age-associated decline in barrier function. Remarkably, treating 26 months old mice with the isoDGR-neutralizing antibody restored BBB integrity to near-youthful levels (Fig. 1G). Collectively, these *in-vivo* findings identify isoDGR accumulation as a causal mediator of neurovascular injury and BBB breakdown in both a genetic model of accelerated molecular aging and in natural aging. Moreover, they establish that neutralization of the isoDGR motif can preserve BBB function, underscoring the central role of this protein damage in age-related cerebrovascular dysfunction.

### IsoDGR impairs endothelial barrier function and cell migration via integrin αV

To determine whether isoDGR directly impairs endothelial cell function as suggested by the BBB leakage observed *in-vivo*, we conducted *in-vitro* experiments using cultured brain endothelial monolayers. Murine bEnd.3 brain microvascular endothelial cells were treated with a synthetic isoDGR-containing peptide, and their barrier properties and migratory capacity were evaluated. A transwell permeability assay^17,42^ revealed that isoDGR exposure significantly increased paracellular permeability. Endothelial monolayers treated overnight with 50 µg/mL isoDGR became substantially more permissive to 60-70 kDa dextran tracer flux compared to untreated controls (Fig. 2A), consistent with our *in-vivo* observation that isoDGR triggers endothelial barrier dysfunction. In parallel, we assessed endothelial cell migration using a scratch-wound assay. Control bEnd.3 monolayers progressively closed the scratch wound over ∼48 hours via collective cell migration. In contrast, isoDGR-treated monolayers showed markedly impaired wound closure, the denuded area remained largely open even after 48 hours, indicating significantly reduced cell motility (Fig. 2B,C). Thus, isoDGR compromises two key endothelial functions, barrier integrity and migration.

**Figure 2.**
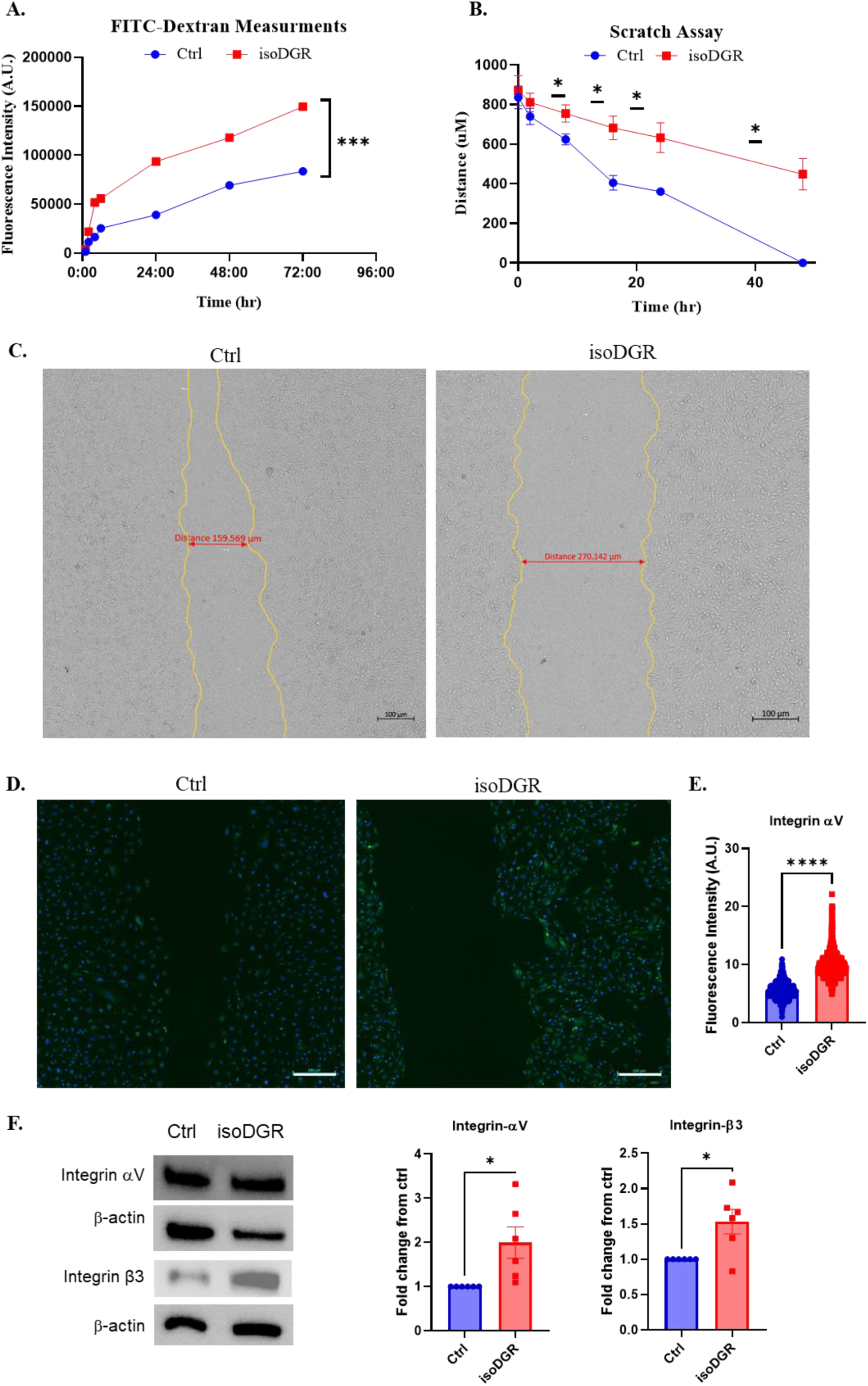
IsoDGR increases permeability, suppresses motility, and induces integrin αV upregulation in brain endothelial cells. (A) Endothelial permeability assay: Confluent monolayers of bEnd.3 brain endothelial cells cultured on Transwell inserts were treated with isoDGR-peptide (50 μg/mL) or vehicle control. Permeability to 60-70 kDa FITC-dextran was measured in the lower chamber at indicated time points. IsoDGR significantly increased paracellular tracer flux, indicating compromised barrier integrity (n = 6 wells per group). Fluorescence was quantified using a fluorometer, and statistical analysis was performed using repeated measures ANOVA. (B) Scratch-wound motility assay: bEnd.3 monolayers were scratched with a pipette tip and cultured in the presence or absence of isoDGR-peptide (50 μg/mL). Wound width was imaged and measured up to 48 hours. IsoDGR treatment significantly delayed wound closure, demonstrating impaired endothelial cell migration (n = 6). Statistical analysis: one-way repeated measures ANOVA. (C) Representative brightfield images of scratched bEnd.3 cultures at 24 hours post-injury. Untreated control cells exhibit narrower wound gaps, whereas isoDGR-treated cells show substantially delayed closure. Red scale bars and yellow outlines demarcate wound width (μm). (D) Integrin αV immunostaining: bEnd.3 cells treated 24 hours with isoDGR-peptide (50 μg/mL) or vehicle control were fixed and stained for Integrin αV (green) and DAPI (blue). Fluorescence imaging revealed enhanced integrin αV signal in isoDGR-treated cells. (E) Quantification of Integrin αV fluorescence intensity across image using Image J. IsoDGR significantly upregulated integrin αV expression (****p < 0.0001, unpaired t-test), suggesting possible involvement of integrin signaling. (F) Densitometric analysis of six independent blots, normalized to control. IsoDGR increases the expression of integrin αV and β3 proteins, consistent with the immunofluorescence results (mean ± SEM; *p < 0.05, unpaired t-test). Scale bars: 100 μm. Error bars represent SEM. These findings support a functional link between isoDGR exposure and endothelial dysfunction via increased permeability, reduced motility, and integrin-mediated signaling.

Because isoDGR structurally mimics RGD motifs that bind integrins, we next examined whether integrin changes accompanied these functional deficits. Indeed, isoDGR-treated endothelial cells displayed an upregulation of integrin αV (ITGAV), which pairs with β3 subunits and normally recognizes RGD sequences. After 24 hours of isoDGR exposure, immunofluorescence staining revealed that integrin αV levels were significantly higher than in untreated cells (Fig. 2D,E). This upregulation was further confirmed by Western blot analysis, which showed increased protein expression of both ITGAV and its binding partner ITGB3 (Fig. 2F). This effect was specific to the intact isoDGR motif, as a control isoD-peptide but lacking the full isoDGR sequence did not trigger any increase in αV. These data suggest that isoDGR engagement of RGD-binding integrins induces a feed-forward increase in integrin expression on endothelial cells. Functionally, increased integrin αV on the cell surface would be expected to strengthen cell-ECM adhesion and stabilize focal adhesion complexes. Paradoxically, excessively strong adhesion can impede cell migration by “locking” cells to their substrate. This appears a plausible explanation for the impaired wound closure: isoDGR-treated cells become hyper-adherent to the matrix and thus less able to move. Additionally, abnormal integrin signaling can weaken cell-cell contacts, for example by altering actin cytoskeletal tension at junctions or downregulating tight junction components. In support of this notion, pathological engagement of endothelial integrin αV/β3 by an RGD-containing protein has been shown increase permeability. It is therefore likely that isoDGR’s effects on barrier function and motility are mediated through integrin αV and its downstream signaling. To gain unbiased mechanistic insights into how isoDGR perturbs endothelial homeostasis, we performed transcriptomic profiling (RNA-seq) of bEnd.3 cells treated with isoDGR versus control.

### IsoDGR reprograms the endothelial transcriptome, disrupting cytoskeletal and junctional networks

Global transcriptomic analysis revealed that isoDGR induces broad gene expression changes in brain endothelial cells. RNA-seq of bEnd.3 cells treated with 50 µg/mL isoDGR for 24 hours (versus controls) identified 515 differentially expressed genes (adjusted p<0.05, |log_2 fold-change| ≥1). Among these, 134 transcripts were upregulated and 381 were downregulated by isoDGR (Fig. 3A). Gene ontology (GO) enrichment analysis indicated that cytoskeletal organization and cell-cell junction pathways were prominently perturbed by isoDGR. Notably, biological processes related to the actin cytoskeleton and cell adhesion were highly enriched among the altered genes, including terms such as intermediate filament cytoskeleton organization, actin crosslink formation, stress fiber assembly, and cell-cell adhesion (Fig. 3B). These transcriptional changes align with our functional observations of impaired barrier integrity and migration, suggesting that isoDGR destabilizes endothelial structure at the genetic program level. Many cytoskelatal-associated and junctional regulators were dysregulated, which likely contributes to the disorganized F-actin architecture and compromised tight junctions observed in isoDGR-treated cells (see below). Consistently, GO categories for cellular components highlighted significant changes in genes associated with the actin cytoskeleton and cell junction (Fig. 3C), while molecular function categories showed enrichment of genes encoding actin-binding and cytoskeletal protein binding activities (Fig. 3D). These data indicate that isoDGR exposure profoundly alters the transcriptional networks governing endothelial cytoskeletal structure and intercellular junction integrity, providing a blueprint for the ensuing barrier dysfunction.

**Figure 3.**
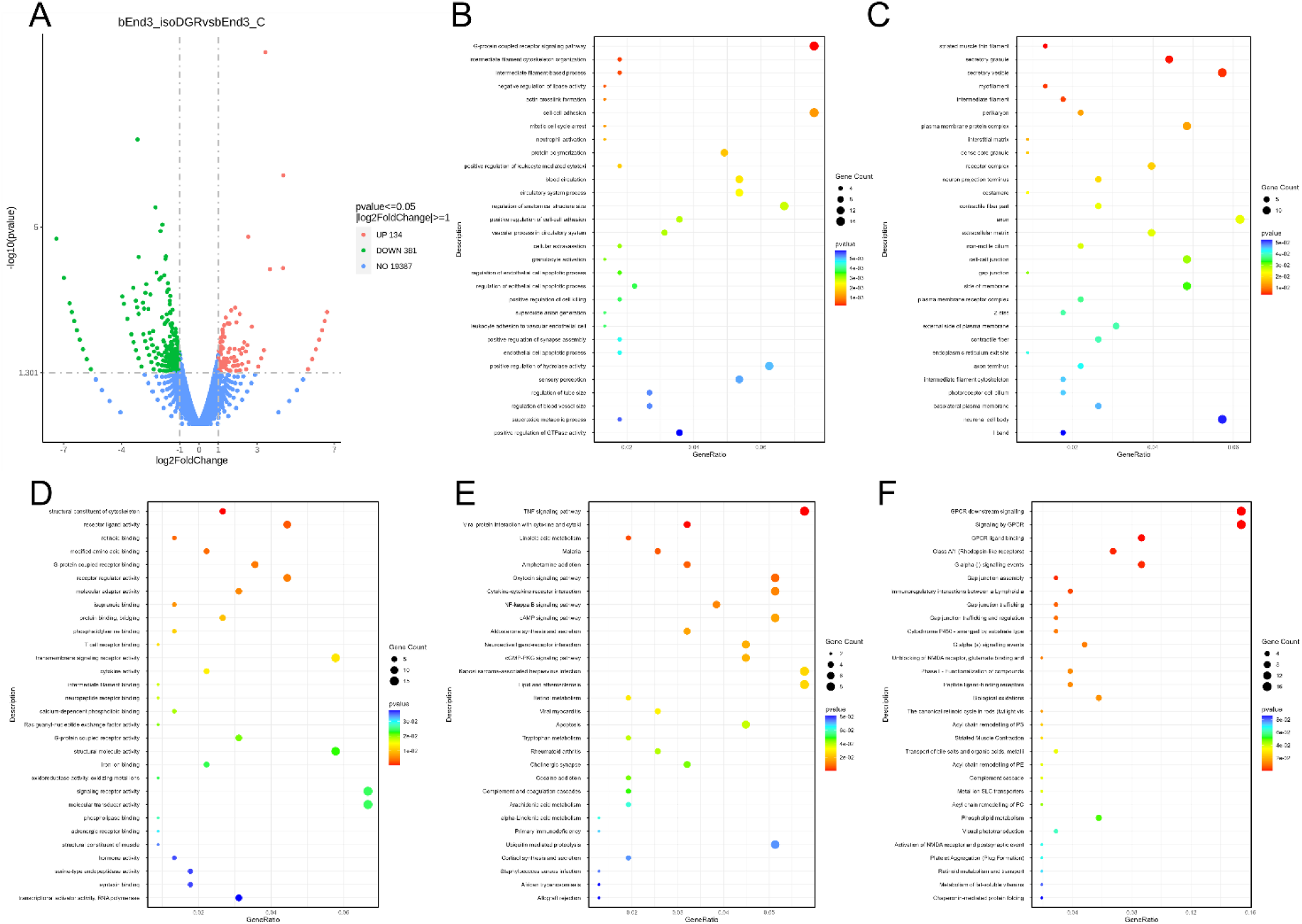
RNA-seq analysis reveals isoDGR-induced cytoskeletal remodeling, junctional disassembly, and endothelial inflammatory priming. *(A)* Volcano plot of differentially expressed genes (DEGs) in bEnd.3 brain endothelial cells treated with 50 μg/mL isoDGR-peptide for 24 hours compared to vehicle-treated controls. A total of 134 genes were significantly upregulated and 381 downregulated (adjusted p < 0.05, |log₂FoldChange| ≥ 1). IsoDGR profoundly alters the endothelial transcriptome, suggesting broad impacts on cellular function and barrier maintenance. (B-D) Gene Ontology (GO) enrichment analysis of differentially expressed genes reveals significant enrichment in pathways related to actin cytoskeleton organization, cell-cell junction assembly, intermediate filament organization, and adherens/tight junction architecture. These changes suggest destabilization of cytoskeletal anchoring and intercellular connectivity, key features of endothelial barrier breakdown. (E-G) Pathway enrichment analysis (KEGG and Reactome) highlights upregulation of inflammatory and stress-related pathways, including TNF signaling, cytokine-cytokine receptor interaction, oxidative stress response, and apoptosis. These shifts reflect endothelial activation, redox imbalance, and pro-death signaling cascades, further compromising vascular function. Together, these transcriptomic data demonstrate that isoDGR drives endothelial dysfunction via cytoskeletal disruption and pro-inflammatory remodeling, the central mechanisms that underlie age-related blood-brain barrier pathology.

In addition to structural pathways, isoDGR triggered a pro-inflammatory transcriptional response. GO biological process terms related to immune cell adhesion and activation were significantly upregulated, including leukocyte migration, granulocyte extravasation, neutrophil activation, and cytokine-cytokine receptor interaction (Fig. 3E). This suggests that isoDGR primes endothelial cells for an inflammatory phenotype conducive to leukocyte recruitment. Pathways associated with oxidative stress and cell fate were also enriched among isoDGR-regulated genes. For example, transcripts involved in superoxide anion generation, reactive oxygen species metabolic process, mitotic cell cycle arrest, and apoptotic signaling were altered, indicating that isoDGR induces oxidative stress and pushes cells toward growth arrest or senescence. Consistent with these patterns, KEGG pathway analysis revealed upregulation of pathways such as TNF signaling and NF-κB signaling, and Reactome analysis identified enrichment of interleukin signaling and other inflammatory networks (Fig. 3F,G). Notably, Ccl2 (also known as MCP-1), a chemokine that recruits monocytes and promotes vascular inflammation, was significantly upregulated in isoDGR-treated endothelial cells. This finding aligns with our previous observations in isoDGR-stimulated immune cells, suggesting that Ccl2 may serve as a shared pro-inflammatory mediator of isoDGR action across vascular and immune compartments. Taken together, The RNA-seq findings delineate a mechanistic framework wherein exposure to isoDGR drives endothelial cells toward a dysfunctional phenotype, characterized by the dysregulation of cytoskeletal and junctional gene networks, potentially compromising barrier integrity, alongside the upregulation of inflammatory and stress-response pathways. These transcriptomic insights guided our next experiments, where we directly examined the structural and inflammatory changes in isoDGR-treated endothelial cells.

### IsoDGR disrupts the endothelial cytoskeleton and intercellular junctions

Our transcriptomic findings suggest that isoDGR has marked effects on cytoskeletal structure. Endothelial barrier integrity depends on a well-organized actin cytoskeleton and intact intercellular junctions to maintain cell-cell contacts. We therefore visualized F-actin fibers and tight junction organization in isoDGR-treated endothelial monolayers via fluorescence microscopy. In control bEnd.3 cells, phalloidin staining revealed prominent, long F-actin stress fibers traversing the cells, and the tight junction protein ZO-1 was arranged in even distribution (Fig. 4A,B, Ctrl). By contrast, isoDGR-treated cells showed a striking disruption of F-actin and ZO-1 organization (Fig. 4A,B, isoDGR). The actin filaments appeared markedly shorter, fragmented, and often collapsed into dense foci or aggregates. ZO-1 localization was also dramatically altered, instead of delineating continuously in cells, ZO-1 in isoDGR-exposed cells was reduced to punctate clusters that were mislocalized away from the distribution in control cells, frequently coalescing with clumped actin. Correspondingly, the normal distribution in control cells was replaced by a jagged and discontinuous pattern in isoDGR-treated monolayers, reflecting compromised cell-cell contacts. To objectively characterize these changes, we performed automated image analysis using the Stress Fiber Extractor (SFEX) algorithm (Fig. S1)^43^. Quantitative analysis revealed that isoDGR treatment led to a significant reduction in total F-actin fiber length, increased fiber fragmentation (decreased mean fiber length and width), and altered fiber orientation compared to controls. Furthermore, isoDGR-treated cells displayed increased cell circularity and decreased cell area, consistent with cytoskeletal collapse and cellular contraction. These findings confirm that isoDGR profoundly disrupts actin stress fiber architecture and cell morphology, as quantified in bEnd.3 monolayers (see Fig. S1).

**Figure 4.**
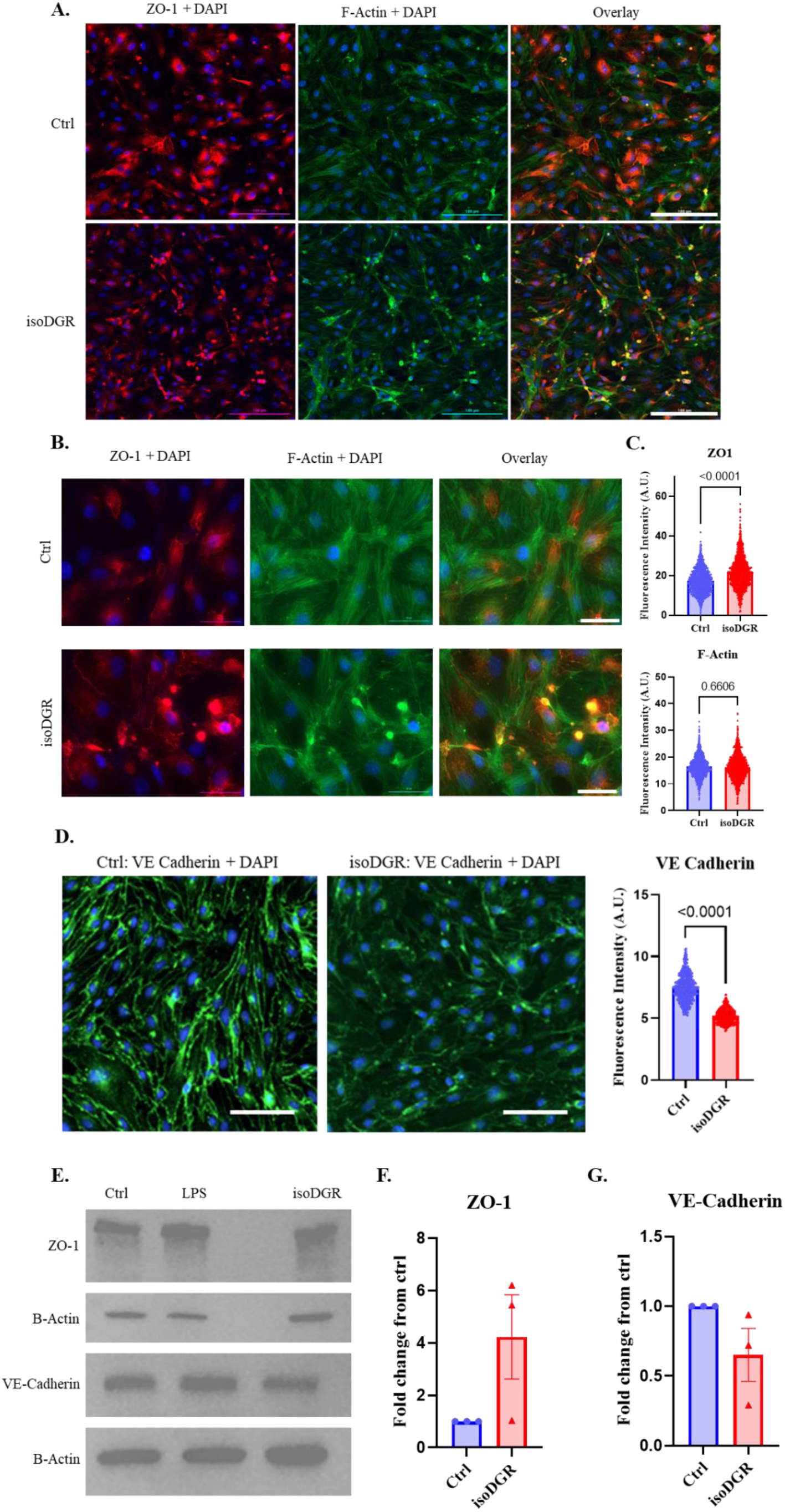
IsoDGR disrupts the cortical actin cytoskeleton and junctional protein organization in brain endothelial monolayers. (A) Low-magnification fluorescence images of confluent bEnd.3 cell monolayers stained for F-actin (phalloidin, green), ZO-1 (red), and nuclei (DAPI, blue) following 24-hour treatment with isoDGR-peptide (50 μg/mL) or vehicle control. IsoDGR exposure causes actin stress fibers and ZO-1 to collapse into aggregated foci. Scale bar: 100 μm. Higher-magnification images (scale bar: 30 μm) illustrate isoDGR-induced collapse of actin filaments and fragmentation of ZO-1 from continuous strands into puncta. Control cells retain organized actin bundles and continuous ZO-1 at cell borders. (C) Quantification of fluorescence intensity for ZO-1 (top) and F-actin (bottom) from immunostained images (≥3 fields/condition, ∼200 ROIs/group, analyzed using ImageJ). IsoDGR significantly increases ZO-1 signal intensity, while F-actin levels remain unchanged, indicating actin redistribution rather than loss. Data are mean ± SEM; unpaired t-test. (D) Representative fluorescence images of VE-cadherin (green) in bEnd.3 cells treated with vehicle or isoDGR (50 μg/mL). Quantification of VE-cadherin intensity (≥3 fields/condition, ∼200 ROIs/group, ImageJ) demonstrates reduced signal after isoDGR treatment. Scale bar: 100 μm. (E) Immunoblot analysis of whole-cell lysates shows increased ZO-1 (∼220 kDa) and decreased VE-cadherin (∼120 kDa) after 24 h isoDGR treatment. β-actin (∼42 kDa) serves as a loading control. LPS (100 ng/mL, 6 h) is shown as a positive control. (F, G) Densitometric quantification from three independent blots, normalized to control. IsoDGR elevates ZO-1 and reduces VE-cadherin protein expression, consistent with immunofluorescence findings (mean ± SEM; unpaired t-test). These results demonstrate that isoDGR exposure destabilizes the endothelial cytoskeleton and disrupts adherens/tight junctions, representing early events in isoDGR-induced blood-brain barrier dysfunction.

We also assessed the status of endothelial adherens junctions by examining vascular endothelial (VE)-cadherin, a key adherens junction protein that maintains cell-cell adhesion. Immunofluorescence staining revealed a pronounced loss of VE-cadherin signal intensity at cell junctions in isoDGR-treated cells compared to controls (Fig. 4D), as quantified by image analysis (Fig. 4D, right panel). Western blot analysis showed that isoDGR exposure led to a consistent decrease in VE-cadherin protein levels (∼30-50% reduction compared to controls, across multiple experiments) (Fig. 4E,G). Loss of VE-cadherin is indicative of junctional destabilization and likely contributes to the increased paracellular permeability observed. Intriguingly, the tight junction protein ZO-1 was found to be upregulated in isoDGR-exposed cells, as evidenced by both immunofluorescence intensity and Western blot band density (Fig. 4C-F). Despite this increase in total ZO-1 protein, its disordered localization (accumulating in cytosolic puncta) suggests a loss of normal tight junction function. We interpret this as a futile compensatory response, isoDGR-triggered signaling induces the endothelial cells to produce more ZO-1, but simultaneously disrupts the actin cytoskeletal scaffold required for ZO-1 functions. The net result is a breakdown of tight junction integrity despite elevated ZO-1 levels. The results indicate that isoDGR disrupts endothelial architecture by collapsing F-actin fibers, fragmenting tight junctions, and reducing adherens junctions. These changes likely underlie the increased permeability and BBB leakage observed with isoDGR. Given that cytoskeletal disruption can trigger inflammation, we next examined whether isoDGR-induced structural changes are linked to inflammatory responses in endothelial cells.

### IsoDGR induces a pro-inflammatory endothelial phenotype

Endothelial cells exposed to isoDGR rapidly adopted an activated, pro-inflammatory phenotype. After 24 hours of isoDGR treatment, bEnd.3 endothelial cells exhibited a marked upregulation of the leukocyte adhesion molecules ICAM-1 and VCAM-1. Immunofluorescence staining revealed approximately a twofold increase in fluorescence intensity of both ICAM-1 and VCAM-1 in isoDGR-treated monolayers compared to untreated controls. In contrast, treatment with the isoD control peptide lacking the isoDGR motif did not significantly alter the expression levels of these adhesion molecules (Fig. 5A-C). Consistently, Western blotting confirmed upregulation of ICAM-1 and VCAM-1 protein in isoDGR-treated cells relative to controls (Fig. 5D). By comparison, the classical inflammatory stimulus lipopolysaccharide (LPS, 100 ng/mL) in our system induced a different adhesion molecule profile, LPS triggered a weaker ICAM-1 increase but a stronger VCAM-1 increase than isoDGR. The pattern elicited by isoDGR (preferential ICAM-1 induction with moderate VCAM-1) points toward activation of an AP-1-dominated signaling pathway, rather than the typical NF-κB-dominant pathway used by LPS. This interpretation is supported by prior evidence that isoDGR engagement can activate ERK-AP1 signaling in vascular endothelial cells^17^.

**Figure 5.**
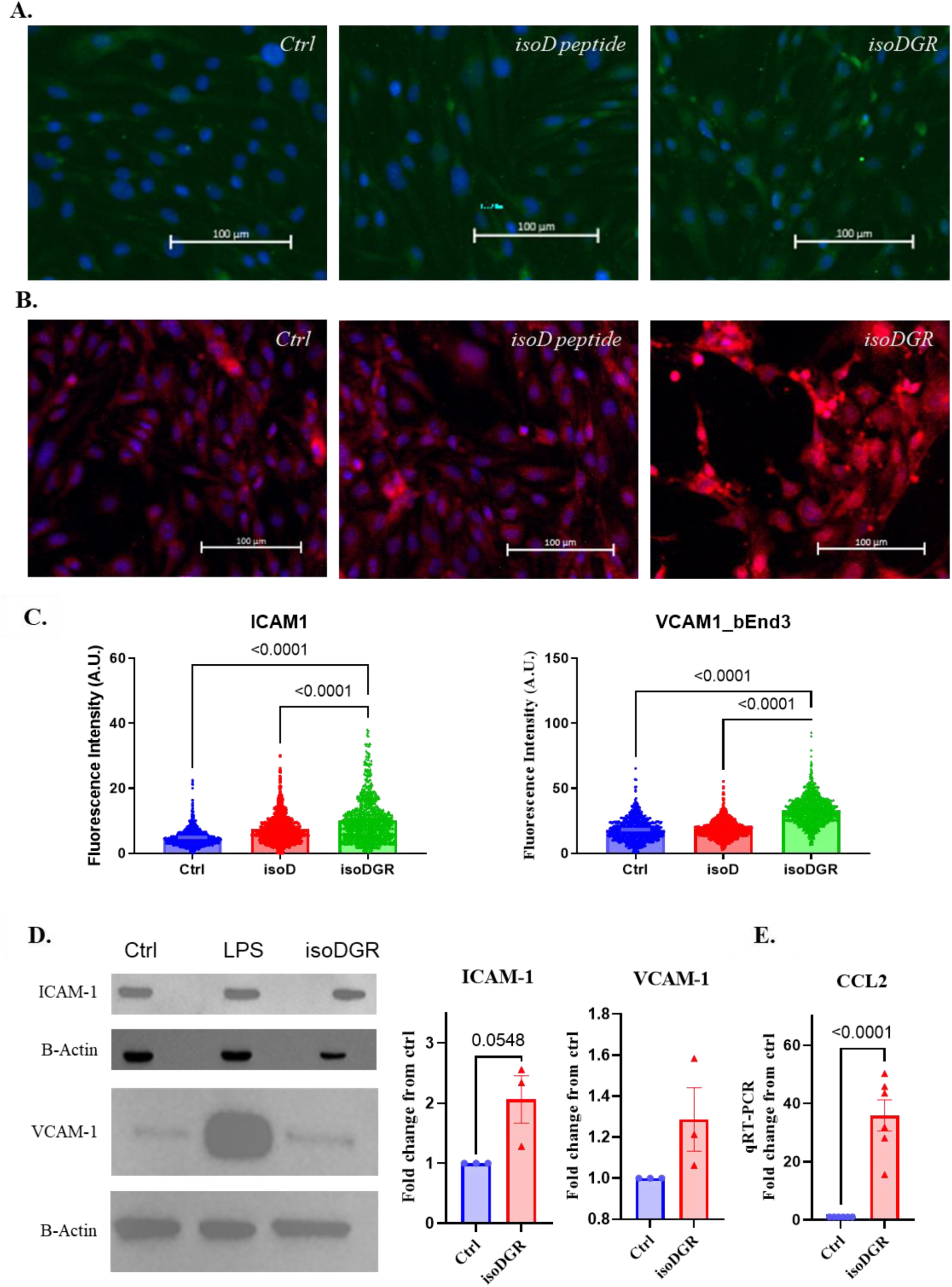
IsoDGR primes brain endothelial cells for leukocyte recruitment by up-regulating ICAM-1, VCAM-1 and CCL2. (A) Immunofluorescence images of bEnd.3 monolayers after 24 h incubation with isoDGR-peptide (50 μg mL⁻¹), an isoaspartate-only control peptide (IsoD, 100 μg mL⁻¹), or vehicle (Ctrl). IsoDGR robustly elevates ICAM-1 surface staining (green); nuclei are counter-stained with DAPI (blue). Scale bar = 100 µm. (B) Parallel staining for VCAM-1 (red) shows a significant increase in isoDGR-treated cells relative to Ctrl and IsoD-peptide. Scale bar = 100 µm. (C) Bar graph quantification of fluorescence intensity (≥3 fields, >200 ROI per condition). IsoDGR induces a ∼2-fold rise in ICAM-1 and VCAM-1, mean ± SEM; p-value from One-way Anova. (D) Immunoblot validation: isoDGR markedly increases ICAM-1 (∼75 kDa) and modestly increases VCAM-1 (∼100 kDa) protein; β-actin serves as loading control. LPS (100 ng mL⁻¹, 6 h) is included as a canonical NF-κB-driven inflammatory control and yields a stronger VCAM-1 than ICAM-1 response. Bar graphs show densitometric means ± SEM from three independent blots. E) qRT-PCR analysis demonstrates significant upregulation of CCL2 (MCP-1) mRNA in isoDGR-treated bEnd.3 cells compared to controls (mean ± SEM; p-values by Student’s t-test). These data are consistent with the transcriptomic signature (Fig. 3) and support the concept that isoDGR activates an ERK-AP-1-biased programme that preferentially drives ICAM-1, VCAM-1 and CCL2/MCP-1 expression, thereby priming endothelial cells for monocyte adhesion and transmigration during isoDGR-mediated BBB dysfunction.

Mechanistically, both cytoskeletal disruption and integrin engagement are known to initiate intracellular pathways that converge on inflammatory gene expression. Integrin “outside-in” signaling through focal adhesion kinases and Rho-family GTPases can lead to ROS production and activation of redox-sensitive transcription factors like AP-1. In our experiments isoDGR exposure induced significant reactive oxygen species generation in endothelial cells (see below), consistent with oxidative stress functioning as a second messenger to drive adhesion molecule upregulation. Notably, the pro-inflammatory changes we observed occurred in the absence of any exogenous cytokines, indicating that the isoDGR-integrin interaction alone is sufficient to intrinsically switch endothelial cells into an activated state including upregulation of CCL2/MCP-1 (as shown by transcriptomic analysis and confirmed by qRT-PCR; Fig. 5E). In such a state, the endothelium would be primed to capture circulating leukocytes, which aligns with our previous *in-vivo* observation that CD68+ monocytes clustered around isoDGR-rich vessels in Pcmt1^−/-^ and aged WT mice^17–19^. These results demonstrate that isoDGR converts endothelial cells into an inflammatory platform, characterized by high expression of leukocyte adhesion molecules and likely facilitating immune cell transmigration. We next examined whether this isoDGR-induced inflammatory activation is associated with oxidative stress, a common mediator of endothelial dysfunction.

### IsoDGR triggers oxidative stress and metabolic dysfunction

Our transcriptomic analysis indicated that isoDGR upregulates pathways related to oxidative stress, suggesting that excessive reactive oxygen species (ROS) might be generated. Many forms of endothelial stress, inflammation, etc. lead to excess ROS production, which can damage cellular components and activate redox-sensitive pathways. To directly test if isoDGR causes oxidative stress, we measured ROS levels in endothelial cells after isoDGR exposure. Using the CellROX Green reagent (which fluoresces upon oxidation by ROS), we found that bEnd.3 cells treated with 50 µg/mL isoDGR-peptide for 24 hours exhibited an increase in cytosolic ROS compared to untreated cells (Fig. 6A). We next assessed mitochondrial ROS production using MitoSOX Green, a fluorogenic indicator of mitochondrial superoxide. Cells exposed to isoDGR showed a marked rise in mitochondrial superoxide levels, especially at higher isoDGR doses and longer exposure times. In a 4-day treatment, 100 µg/mL isoDGR caused strong MitoSOX fluorescence, comparable to the positive control (0.2 mM doxorubicin), indicating significant mitochondrial ROS accumulation (Fig. 6B). Quantification of the fluorescence signals confirmed that isoDGR treatment significantly elevated both total cellular ROS and mitochondrial ROS in a dose- and time-dependent manner (Fig. 6C,D). Using an alternate ROS assay (DCFH-DA on cell lysates)^44^, we observed similarly that isoDGR exposure increased endothelial ROS production ∼1.5-fold relative to controls after 4 days (Fig. 6E). These data demonstrate that isoDGR not only stimulates cytosolic ROS generation but also perturbs mitochondrial function, leading to leakage of reactive oxygen from the electron transport chain.

**Figure 6.**
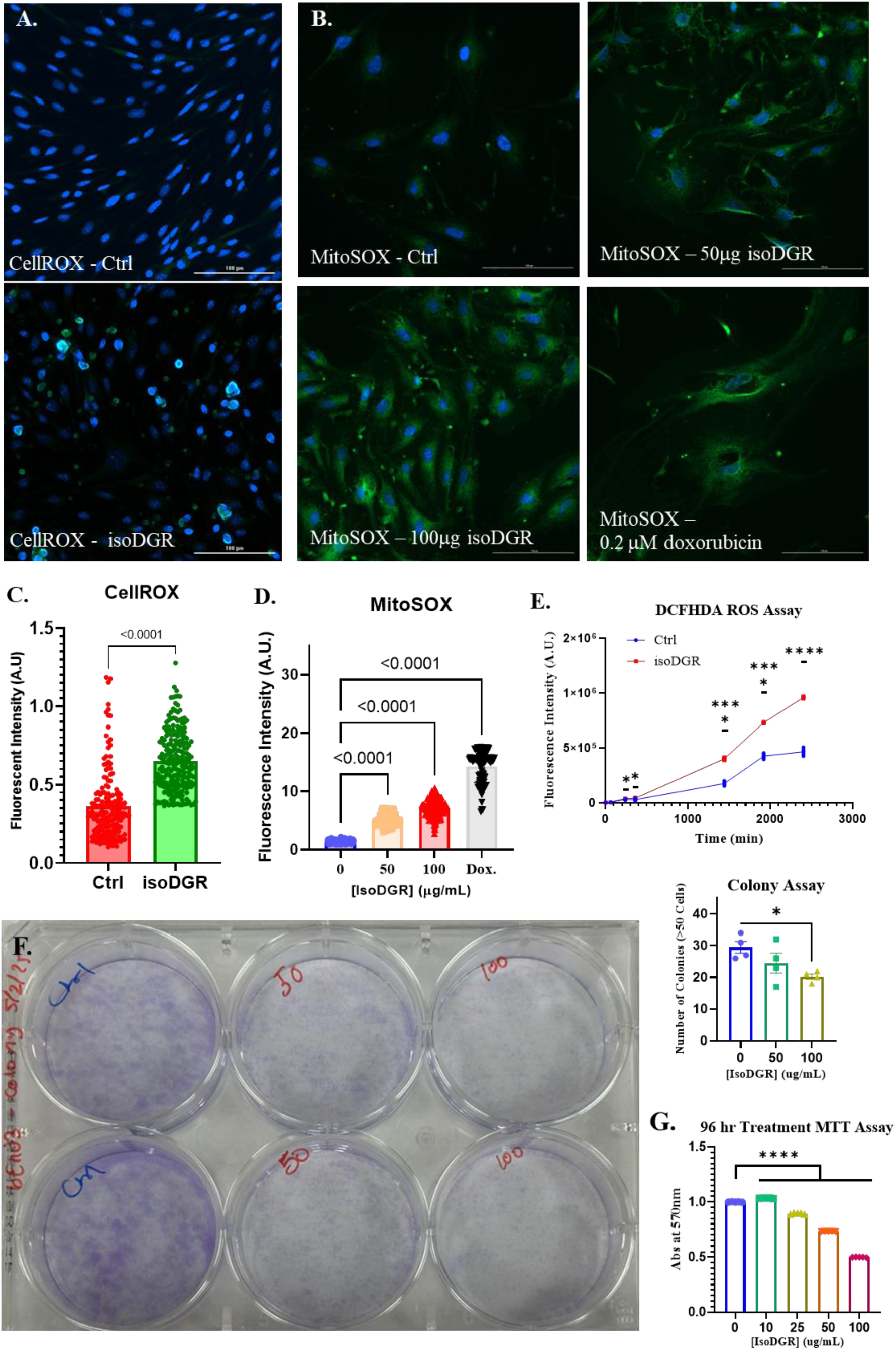
IsoDGR provokes oxidative stress and suppresses proliferative capacity in brain endothelial cells in key upstream events in isoDGR-driven endothelial senescence. (A) Cytosolic ROS: bEnd.3 monolayers were exposed to isoDGR-peptide (50 µg mL⁻¹, 24 h) or vehicle (Ctrl), then incubated with CellROX Green (5 µM, 30 min, 37 °C). IsoDGR markedly increases punctate CellROX fluorescence (oxidized ROS sensor). Scale bar = 100 µm. (B) Mitochondrial superoxide: Cells were treated with isoDGR (0, 50, 100 µg mL⁻¹) or doxorubicin (0.2 µM, positive ROS control) and stained with MitoSOX™ Green (30 min). Dose-dependent mitochondrial ROS accumulation is evident; doxorubicin shows comparable intensity. Scale bar = 100 µm. (C) Bar graph quantification of CellROX fluorescence (ImageJ, ≥3 fields per condition). IsoDGR increases cytosolic ROS ∼2-fold (mean ± SEM; unpaired t-test). (D) Bar graph quantification of MitoSOX fluorescence showing significant elevations with 50 µg and 100 µg isoDGR and with doxorubicin as positive control (one-way ANOVA). (E) Kinetics of total intracellular ROS measured by DCFH-DA in cell lysates from control vs. isoDGR-treated cultures (50 µg mL⁻¹, 24 h). IsoDGR elevates ROS over post-lysis time course (n = 5, repeated-measures ANOVA). (F) Colony-formation assay: Crystal-violet-stained colonies after 7 days reveal dose-dependent suppression of clonogenic growth by isoDGR (50 and 100 µg mL⁻¹). Bar graph quantifies colonies per 100 000-pixel² ROI (mean ± SEM; * p < 0.05, one-way ANOVA). (G) MTT viability assay (96 h): Metabolic activity declined in a concentration-dependent manner with isoDGR (n = 5; **** p < 0.0001, one-way ANOVA with multiple comparisons). Together, these data show that isoDGR generates both cytosolic and mitochondrial ROS, impairs endothelial metabolic activity, and limits proliferative potential, molecular conditions that precede the senescent phenotype described in Figure 7 and underpin isoDGR-mediated BBB dysfunction.

Excess ROS damage proteins, lipids, and DNA, and can also act as a signaling molecule that modifies cell behavior. In endothelial cells, ROS are known to disrupt tight junction integrity and actin dynamics, thereby contributing to BBB breakdown. Consistent with a ROS-mediated mechanism, our finding that isoDGR disrupts F-actin/ZO-1 organization may be partly explained by oxidative stress, as ROS can cause actin depolymerization and junctional protein dysfunction. Additionally, ROS can activate transcription factors such as NF-κB, which would complement integrin-initiated AP-1 signaling in driving ICAM-1/VCAM-1/MCP-1 expression.

We next evaluated whether isoDGR impacts endothelial cell proliferation and viability, as persistent oxidative stress is often antiproliferative. A 7-day cell growth assay revealed that isoDGR indeed exerted anti-proliferative effects. bEnd.3 cells cultured in the presence of isoDGR formed fewer and smaller colonies over a week compared to untreated controls, in a dose-dependent manner (with 100 µg/mL isoDGR significantly reducing colony numbers; Fig. 6F). Similarly, metabolic activity measured by the MTT assay showed that isoDGR caused a concentration- and time-dependent decline in endothelial cell viability. At 50 µg/mL isoDGR, MTT values were reduced by ∼30% over 4 days relative to control cultures (Fig. 6G). Microscopic inspection of the cells corroborated a lower cell density and morphological changes consistent with cell cycle arrest or senescence (enlarged, flattened cells), rather than acute cell death. Notably, the timing of these anti-proliferative effects coincided with the emergence of high ROS levels in isoDGR-treated cells, implicating oxidative stress as a key mediator of growth arrest. Mitochondrial dysfunction caused by isoDGR could also impair ATP production and alter redox homeostasis, potentially depleting NAD+ and triggering DNA damage responses that enforce a cell-cycle arrest. The reduction in MTT (which depends on mitochondrial NAD(P)H production) further supports a loss of metabolic competence in isoDGR-exposed cells. Collectively, these data indicate that isoDGR creates a state of oxidative stress and metabolic dysfunction in endothelial cells, which impairs their proliferative capacity and hints at an impending transition to a senescent phenotype. We therefore examined classical markers of cellular senescence to determine if prolonged isoDGR exposure indeed drives endothelial cells into premature senescence.

### IsoDGR drives endothelial cells into premature senescence

After 4 days of continuous isoDGR exposure, brain endothelial cells exhibited distinct signs of premature senescence. IsoDGR-treated bEnd.3 cells showed a marked increase in senescence-associated β-galactosidase (SA-β-gal) activity compared to untreated controls. Quantitatively, the percentage of SA-β-gal-positive cells was significantly increase by isoDGR treatment (Fig. 7A,D,H). IsoDGR-treated cells also displayed the enlarged, flattened morphology characteristic of senescent cells. In contrast, cells treated with a control isoD-peptide without the isoDGR-motif showed minimal SA-β-gal staining, indicating that the complete isoDGR motif is specifically responsible for driving endothelial cells into senescence beyond any non-specific stress from isoAsp residues.

**Figure 7.**
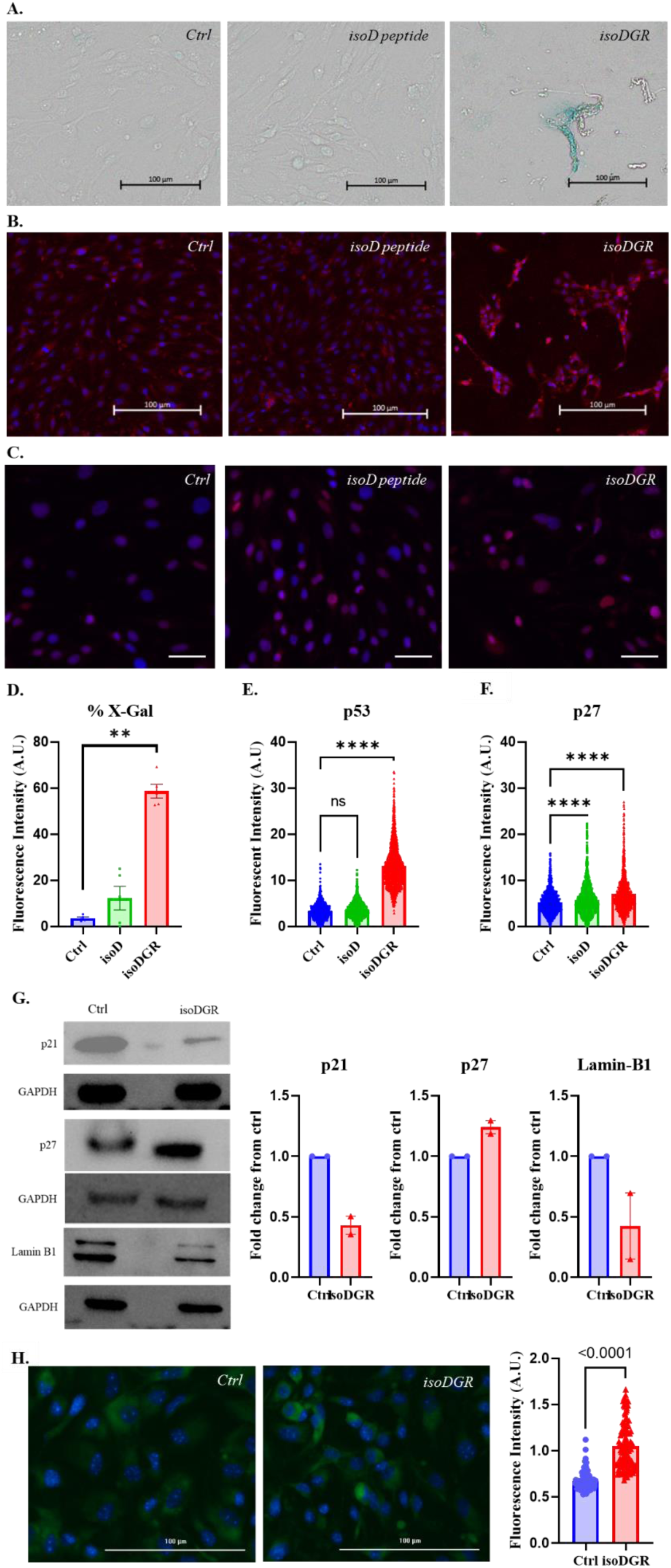
Sustained isoDGR exposure drives brain endothelial cells into an atypical, p53/p27-biased senescent state, an end-stage of the isoDGR injury cascade. (A) Four-day exposure of bEnd.3 monolayers to isoDGR-peptide (50 µg mL⁻¹) triggers widespread senescence-associated β-galactosidase (SA-β-gal) staining (teal) compared with vehicle-treated (Ctrl) or iso-Asp control peptide (IsoD) cultures. Scale bars = 100 µm. (B, C) Immunofluorescence for the senescence regulators p53 and p27 (red) shows strong nuclear accumulation in isoDGR-treated cells, whereas Ctrl and IsoD-peptide cause only basal staining. Scale bars = 100 µm; right-hand panels show higher magnification. (D) Quantification of SA-β-gal-positive cells per 100 000-pixel² ROI confirms a robust increase after isoDGR (mean ± SEM; **p < 0.01, ****p < 0.0001, n = 5, one way ANOVA). (E, F) Bar graphs of p53 and p27 fluorescence intensities demonstrate significant up-regulation in isoDGR-treated monolayers (mean ± SEM; ****p < 0.0001, one way ANOVA). (G) Immunoblot analysis corroborates the atypical senescence signature: p27 proteins rise, lamin B1 decreases, whereas p21 shows decrease. GAPDH, loading control. Bar graphs summarise two independent blots. (H) Live-cell Spider-β-gal reporter (green) confirms elevated senescence signal in isoDGR-treated cultures; Bar graph quantifies fluorescence (mean ± SEM; unpaired t-test, n = 5). Scale bar = 50 µm. Together with the oxidative-stress and growth-inhibition data (Fig. 6), these results place endothelial senescence as a downstream consequence of isoDGR-induced cytoskeletal collapse, ROS generation, and inflammatory activation, providing a mechanistic endpoint that links ECM damage to irreversible BBB dysfunction in ageing vasculature.

IsoDGR exposure also modulated key cell cycle regulators associated with the senescent state. Endothelial cells conditioned with isoDGR exhibited substantially higher levels (∼2-fold) of the tumor suppressor protein p53, as visualized by immunostaining (Fig. 7B,E). Surprisingly, the cyclin-dependent kinase inhibitor p21 (a canonical downstream effector of p53 in the senescence pathway) was not upregulated in isoDGR-treated cells; in fact, p21 levels were reduced relative to controls as revealed by WB (Fig. 7G). This suggests that isoDGR triggers a senescence program that does not primarily rely on the classic p53-p21 axis. Instead, another CDK inhibitor, p27^Kip1^, was found to be upregulated. IsoDGR-treated cells showed elevated nuclear p27 by immunofluorescence (Fig. 7C) and an increase in total p27 protein by WB (Fig. 7G). In addition, the nuclear envelope protein lamin B1, which is typically downregulated during senescence, was significantly diminished in isoDGR-exposed cells (to ∼50% of control levels, Fig. 7G). This combination of changes (high p53 and p27, low lamin B1, with little change in p21) strongly indicates that chronic isoDGR exposure induces a senescent phenotype via an atypical pathway in chronic isoDGR exposure.

Taken together, our results demonstrate that isoDGR acts as a potent inducer of premature endothelial senescence, characterized by permanent growth arrest and a pro-inflammatory, adhesion-molecule-rich state. Senescent endothelial cells, in turn, are known to exacerbate vascular pathology and by failing to maintain or repair the BBB. Notably, the chain of events we observed, from acute isoDGR effects (integrin hyperactivation and cytoskeletal/junctional disruption) to chronic outcomes (inflammation and senescence), provides a coherent timeline of how age-related ECM modifications may drive endothelial aging. In essence, isoDGR accumulation initiates a cascade of endothelial dysfunction that begins with cytoskeletal collapse and barrier breakdown and transforms in a senescent, pro-inflammatory endothelial phenotype.

## DISCUSSION

In this study, we identify isoDGR as an aging-associated ECM modification that drives endothelial dysfunction and BBB breakdown. Our data from mouse models and cell culture assays reveal how the accumulation of this single matrix lesion can precipitate in neurovascular pathology. IsoDGR thereby emerges as a molecular link between vascular aging, chronic inflammation, and BBB-mediated neurodegeneration.

A key insight from our work is that isoDGR acts through integrin receptors to perturb endothelial cell structure and function. In healthy tissues, integrins continuously sense the ECM and help maintain endothelial stability. However, isoDGR-integrin engagement appears to redirect these signals toward deleterious outcomes. We observed an upregulation of integrin αV and β3 in endothelial cells exposed to isoDGR, consistent with prior reports that isoDGR can increase surface integrin expression on vascular cells. This likely creates a feed-forward loop of enhanced integrin clustering and cell-matrix adhesion. As a result, isoDGR-treated endothelial cells become uncoupled from their neighboring cells, as evidenced by the collapse of actin stress fibers and the disruption of intercellular junctions, which led to loss of BBB integrity in our *in-vivo* models. Mechanistically, integrin “outside-in” signaling can activate focal adhesion kinase and myosin light-chain pathways that increase cellular contractility, or conversely cause a loss of cytoskeletal tension; both scenarios can destabilize intercellular junctions. Our finding of actin fiber dissolution and tight junction dissociation aligns with these mechanisms. Notably, similar cytoskeletal and junctional derangements have been observed when endothelial αV/β3 integrins are aberrantly engaged by RGD-containing ligands^45,46^. Oxidative stress may further compound this effect, integrin signaling platforms can recruit NADPH oxidases that generate ROS^47^, and integrin-induced ROS have been shown to disrupt the cytoskeleton and tight junctions in endothelial cells^48^. Thus, isoDGR’s ability to hyper-activate integrins likely initiates a cascade of cytoskeletal collapse and junctional failure, providing a direct path from ECM damage to BBB breakdown.

Another major finding is that isoDGR directly promotes an inflammatory activation of endothelial cells. Classically, chronic endothelial inflammation in aging has been attributed to circulating cytokines or pathogen-sensing receptors (e.g. Toll-like receptors). Our results highlight an alternative paradigm, a spontaneous molecular damage to the ECM in endothelial basement membrane can directly activate endothelial integrins and incite inflammation from within. IsoDGR exposure led to upregulation of ICAM-1, VCAM-1, and the chemokine MCP-1/CCL2 in brain endothelial cells without any external inflammatory stimulus. This finding indicates that isoDGR-integrin interactions can initiate an NF-κB/AP-1 driven gene expression program characteristic of endothelial activation^17^. The specific pattern of adhesion molecule expression we observed implicates AP-1 involvement, aligning with evidence that isoDGR triggers ERK-AP1 signaling in endothelial and monocytic cells^17^. Interestingly, the pro-inflammatory shift induced by isoDGR closely resembled the senescence-associated secretory phenotype (SASP) of endothelial cells, wherein senescent cells overexpress adhesion molecules and cytokines^48–50^. This suggests a mechanistic link between ECM damage and “sterile” inflammation in aging. Together, these data support the concept that isoDGR damage transforms the endothelium into a chronically inflamed state, which likely contributes to BBB breakdown and neuroinflammation in the aging brain.

Oxidative stress appears to be a critical mediator connecting isoDGR to these pathological outcomes. We found that isoDGR markedly increased both cytosolic and mitochondrial ROS in endothelial cells. Excess ROS can damage DNA (activating p53-driven pathways), oxidize signaling proteins, and further deteriorate the ECM. Integrin signaling complexes are known to assemble NADPH oxidase enzymes that produce ROS^47^, providing one route by which isoDGR-integrin engagement elevates oxidative stress. In the brain endothelium, oxidative stress is a well-known mechanism of BBB impairment in conditions ranging from acute stroke to Alzheimer’s disease. Our results suggest that accumulation of isoDGR in aging microvessels may chronically raise endothelial ROS levels, causing the BBB to become leaky even in the absence of overt inflammation. This idea ties in with clinical observations that BBB breakdown is an early event in cognitive aging and can predict progression to dementia^1–3^. By driving sustained ROS production, isoDGR might thus serve as an intrinsic accelerator of BBB aging and neurodegenerative risk.

The most profound consequence of chronic isoDGR exposure we observed was the induction of endothelial senescence. Endothelial cell senescence is increasingly recognized as a contributor to age-related vascular disease, promoting thrombogenicity, barrier permeability, and inflammation ^51–56^. Our findings indicate that isoDGR damage can drive premature endothelial senescence, a novel concept linking ECM aging to cellular aging. We documented multiple classic markers of senescence in isoDGR-treated cells: increased SA-β-gal activity, activation of p53, upregulation of p27^Kip1^, and loss of lamin B1, all indicative of a senescent phenotype. Interestingly, p21^Cip1^ was not elevated, suggesting a noncanonical senescence pathway is at play. Mechanistically, sustained integrin hyper-activation (due to isoDGR), continuous ROS production, and chronic stress signaling (e.g. ERK/AP-1) likely converge to trigger growth arrest and senescence. Indeed, isoDGR-treated endothelial cells ceased proliferating and adopted an inflammatory profile, which would further propagate vascular dysfunction *in-vivo*. Our findings revealed that senescent endothelial cells in the cerebral microcirculation are key drivers of BBB dysfunction in aging. By forcing endothelial cells into a senescent, pro-inflammatory state, isoDGR may thus act as an upstream catalyst of the BBB breakdown and microvascular rarefaction observed in the aged brain.

Overall, our study identifies isoDGR as a critical mediator of age-related vascular inflammation and BBB dysfunction. Prior work in large arteries linked isoDGR formation to monocyte adhesion and atherogenesis; we now extend these insights to the cerebral microvasculature and BBB integrity. It is conceivable that isoDGR accumulation in aging human brains (for example, in long-lived ECM deposits) contributes to the pathogenesis of neurodegenerative diseases. Chronic BBB leakiness and neuroinflammation are early features of vascular dementia, Alzheimer’s disease and related dementias, and if isoDGR is present in aged or AD brains, it could provide a tangible molecular link between vascular aging and neurodegeneration. Encouragingly, recent studies in animal models have shown that targeting isoDGR with specific antibodies can ameliorate age-related pathology^18,19^. In Pcmt1^−/-^ mice, anti-isoDGR immunotherapy reduced systemic inflammation and significantly extended lifespan. In the present study, an isoDGR-neutralizing antibody restored BBB integrity in both aged wild-type and Pcmt1^−/-^ mice, underscoring the therapeutic potential of this approach. Blocking isoDGR-integrin interactions might prevent the cascade of integrin activation, ROS generation, inflammatory signaling, and cellular senescence that culminates in BBB failure. This strategy might be beneficial not only for vascular aging, but also for other chronic conditions involving ECM damage, such as atherosclerosis, in which the isoDGR-laden matrix has been shown to promote immune cell recruitment.

In summary, our work supports a model in which age-related isoDGR modification of the ECM triggers endothelial cytoskeletal disintegration and inflammatory activation, thereby compromising the BBB. These data broaden the prevailing paradigm of vascular aging to include the ECM as an active driver of endothelial pathology. By delineating the pathway from isoDGR accumulation through integrin signaling to oxidative stress, inflammation, and senescence, we provide a framework for targeting these processes *in-vivo*. Therapeutically, intervening in the isoDGR-integrin axis could break the vicious cycle of molecular damage and cellular dysfunction. Our findings, combined with evidence of anti-isoDGR antibody efficacy, suggest that immune clearance or neutralization of this ECM damage motif is a viable strategy to preserve BBB integrity and dampen neuroinflammation during aging. Such interventions, for example, an isoDGR-specific monoclonal antibody, might help maintain cognitive function by protecting the brain’s vasculature as we grow older.

## MATERIALS AND METHODS

### Mouse Models and Antibody Treatment

All animal studies were conducted in accordance with protocols approved by the Nanyang Technological University Institutional Animal Care and Use Committee (IACUC protocols #ARF-SBS/NIE/LKC-A18016, A19029, A18059). Pcmt1 knockout (Pcmt1^−/-^ mice on a C57BL/6J genetic background were used to investigate the effects of unrepaired isoDGR accumulation. These mice were generated by intercrossing Pcmt1^+/-^ heterozygotes, as previously described^18,19,39^, and genotyped by PCR to confirm the absence of the PCMT1 gene. Age-matched wild-type (WT) littermates (Pcmt1^+/+^) were used as controls for all experiments involving Pcmt1^−/-^ mice.

For accelerated isoDGR accumulation studies, cohorts of male Pcmt1^−/-^ and WT littermates, of age 8-10 weeks, were used. For natural aging studies, male C57BL/6J WT mice were obtained from an in-house colony and aged to either 3 months (“young”) or 26 months (“aged”) prior to experimentation.

For antibody intervention studies, mice received intraperitoneal injections of purified isoDGR-specific monoclonal antibody (isoDGR-mAb; 1 mg/kg body weight) once weekly for 8 consecutive weeks^18,19^. Control groups received equal volumes of isotype-matched control IgG (Bio X Cell) on the same schedule. Antibody dosing was initiated 8 weeks prior to the start of experimental procedures. Group sizes ranged from 3 to 8 animals per experimental condition. Mice were randomly assigned to treatment or control groups, and all interventions and outcome assessments were performed in a blinded manner to minimize experimental bias.

Throughout the study, only male mice were used to avoid potential confounding effects of sex. Animals were housed in individually ventilated cages under specific pathogen-free conditions, maintained on a 12-hour light/dark cycle, and provided unrestricted access to autoclaved standard chow and water. Animal health and welfare were monitored daily, and all procedures were performed in accordance with institutional and national guidelines for the ethical use of animals in research.

### Anti-isoDGR Monoclonal Antibody Generation and Administration

The isoDGR-specific monoclonal antibody (isoDGR-mAb) was generated as previously described^17^. In brief, BALB/c mice were immunized with a synthetic isoDGR-containing peptide (GC[isoD]GRCGK) conjugated to a Keyhole Limpet Hemocyanin carrier protein. Hybridoma technology was used to produce monoclonal antibodies by Genscript (USA). The hybridoma supernatants were screened by ELISA for high-affinity and isoform-specific binding to the isoDGR motif, with minimal cross-reactivity to related sequences (isoD, NGR, DGR, or RGD).

For *in-vivo* experiments, purified isoDGR-mAb was administered to mice by intraperitoneal injection at a dose of 1 mg/kg body weight, once weekly, for a total of eight weeks prior to analysis. Control mice received an equivalent dose and schedule of an isotype-matched control monoclonal antibody (mouse IgG, Bio X Cell). All antibody preparations were sterile-filtered and endotoxin levels were confirmed to be below the accepted threshold for animal use.

### Evans Blue Extravasation Assay

To evaluate blood-brain barrier (BBB) integrity, Evans blue dye extravasation was performed as previously described^39^, with minor modifications. Briefly, mice were injected via the tail vein with 100 μL of 4% Evans blue solution (E2129, Sigma-Aldrich) in sterile PBS. The dye was allowed to circulate systemically for 30 minutes. Mice were then deeply anesthetized with isoflurane and perfused intracardially with cold sterile PBS to remove intravascular dye. Brains were rapidly harvested, weighed, and incubated in 1 mL formamide (Sigma-Aldrich) at 45°C for 48 hours to extract Evans blue from the tissue. Absorbance of the supernatant was measured at 650 nm using a Universal Microplate Reader (BioTek Instruments).

### Fluorescent Dextran Extravasation Assay

For quantitative analysis of BBB permeability to high and low molecular weight tracers, mice received a single intravenous injection (tail vein) of either 70 kDa Texas Red-conjugated dextran (100 μg/g body weight; D1830, Thermo Fisher Scientific) or 4 kDa FITC-dextran (10 μg/g; 46944, Sigma-Aldrich), diluted in sterile PBS. For studies involving Pcmt1^−/-^ and age-matched Pcmt1^+/+^ mice (with or without isoDGR-mAb or isotype treatment), 70 kDa Texas Red-dextran was used, while 4 kDa FITC-dextran was used in naturally aged mice to detect subtle barrier disruption. Tracers were allowed to circulate for 2 hours. Subsequently, mice were anesthetized with isoflurane and perfused transcardially with cold PBS for 10 minutes at 4°C to remove intravascular dye. Brains were removed and sectioned (Sagittal slices), and fluorescence images were acquired using an epifluorescence microscope (Nikon Eclipse Ti-2).

Fluorescence intensity within defined brain regions of interest (ROI) was quantified in at least three independent experiments, using ImageJ software. For each condition, at least 200 ROIs were analyzed per experiment to provide quantitative assessment of tracer extravasation and BBB integrity.

### Cell Culture and Reagents

Mouse brain microvascular endothelial cells (bEnd.3, ATCC CRL-2299) were cultured in Dulbecco’s Modified Eagle Medium (DMEM; D7777, Sigma-Aldrich) supplemented with 10% heat-inactivated fetal bovine serum (FBS; ThermoFisher Scientific), 1% Penicillin-Streptomycin (ThermoFisher Scientific), and maintained at 37°C in a humidified incubator with 5% CO₂. Cells were routinely passaged at 70-80% confluence using 0.05% trypsin-EDTA (25300054, ThermoFisher Scientific) and used for experiments between passages 5 and 15. Culture media were replaced every 2-3 days.

IsoDGR-peptide (GC(isoD)GRCGK) and isoD-peptide as control were synthesized (GenScript) and prepared as 10 mg/mL stocks in sterile, endotoxin-free 1× PBS, then diluted to working concentrations (typically 50 μg/mL) immediately before use. All experimental treatments were performed on cells at 60-80% confluency unless otherwise specified. For positive control of senescence or oxidative stress, doxorubicin (44583, Sigma-Aldrich) was freshly prepared in sterile water and used at concentrations indicated in each experiment.

All reagents with their catalog numbers, including antibodies and fluorescent probes, were obtained from manufacturers as indicated in each protocol section below.

### Transwell Endothelial Permeability Assay

Endothelial barrier integrity was assessed using a transwell permeability assay. Briefly, 24-well permeable cell culture inserts with a 0.4 μm pore size (Corning, or A1048301, ThermoFisher Scientific) were coated with rat tail type I collagen (A1048301, ThermoFisher Scientific). Collagen solution was applied evenly to the upper membrane surface, and inserts were air-dried in a sterile culture hood for 30 minutes before use.

bEnd.3 cells were seeded onto the apical (upper) surface of the coated transwell membranes at a density of 1.5 × 10⁵ cells per insert and cultured in DMEM medium as described above. Each insert and the corresponding lower well received 400 μL of complete medium. Cells were incubated at 37°C in 5% CO₂ and allowed to reach 100% confluence (typically 2-3 days). Upon confluency, the culture medium was replaced with fresh medium containing IsoDGR-peptide at a final concentration of 50 μg/mL. Control inserts received medium containing isoD-peptide or without treatment. After 24 hours of treatment, endothelial permeability was assessed by adding 0.2 mg/mL FITC-labeled dextran (59,000-77,000 g/mol; FD70, Sigma-Aldrich) to the upper chamber.

At designated time points, 10 μL of medium was sampled from the lower chamber and immediately diluted in 190 μL of fresh medium. FITC-dextran fluorescence was measured at excitation/emission wavelengths of 480/520 nm using a DeNovix fluorometer. Background fluorescence (medium only) was subtracted. All conditions were analyzed in at least triplicate, and results were normalized to the blank control and expressed as mean ± SEM.

### Scratch Wound Migration Assay

Endothelial cell migration was evaluated using a scratch wound assay. bEnd.3 cells were seeded into 6-well tissue culture plates (Corning) at a density to reach approximately 90% confluency in DMEM medium. The cell monolayer was washed gently with pre-warmed PBS to remove residual serum and debris. A linear wound (scratch) was created in each well using a sterile 100 μL pipette tip. Detached cells were removed by washing the wells twice with PBS. Fresh culture medium (with IsoDGR-/isoD-peptides at 50 μg/mL final concentration or without treatment) was added to the respective treatment and control wells. Plates were immediately returned to the incubator (37°C, 5% CO₂). Wound closure was monitored using a Cytation5 Imaging Reader (BioTek) in brightfield mode at defined time points: 0, 2, 4, 8, 24, and 48 hours after scratching. At each interval, images were captured at standardized locations along the wound. The distance between the wound edges was measured at 10 evenly spaced points per well using ImageJ, and the mean gap width was calculated for each condition at each time point. For immunofluorescence analysis, parallel scratch assays were performed on cells seeded on coverslips, following the same protocol, to allow post-assay fixation and staining. No extracellular matrix coating was applied to coverslips in these assays. All experiments were conducted in at least triplicate. Investigators performing quantification were blinded to experimental group assignments.

### Transcriptomic Analysis (RNA-seq and Pathway Analysis)

The bEnd.3 cells were cultured and treated with isoDGR-peptide (50 μg/mL) or vehicle control for 24 hours. Total RNA was extracted using the RNeasy Mini Kit (Qiagen) according to the manufacturer’s protocol. RNA integrity and concentration were assessed using an Agilent 2100 Bioanalyzer and a NanoDrop spectrophotometer, respectively. RNA samples with RNA integrity number (RIN) >8 were used for library preparation.

RNA-seq libraries were constructed using the NEBNext Ultra II RNA Library Prep Kit (New England Biolabs) and sequenced on a NovaSeq X Plus platform (Novogene Corporation Inc., Sacramento, CA, USA) to generate paired-end 150 bp reads. Quality control was performed using FastQC, and adapter sequences were trimmed with Trimmomatic. Clean reads were mapped to the mouse reference genome (GRCm38) using STAR aligner. Gene-level quantification was performed with featureCounts, and differential gene expression analysis was conducted using DESeq2. Significantly differentially expressed genes were identified using an adjusted p-value < 0.05 and |log2(fold change)| ≥ 1.

Pathway enrichment analysis was performed using the clusterProfiler package to identify overrepresented Gene Ontology (GO) biological processes and Kyoto Encyclopedia of Genes and Genomes (KEGG) pathways. Data visualization included volcano plots and bubble plots to highlight the most enriched pathways and gene categories.

### Immunofluorescence Staining

The bEnd.3 cells were seeded onto sterile glass coverslips placed in 6-well plates and cultured in DMEM medium at 37°C in 5% CO₂. Once cells reached approximately 70% confluence, they were treated with isoDGR-peptide to final concentration of 50 μg/mL for 24-48 hours. Untreated cells served as negative controls. For F-actin staining, cells were incubated with phalloidin-iFluor 488 (ab176753, Abcam) or phalloidin-iFluor 594 (ab176757, Abcam) following the manufacturer’s protocol prior to antibody incubation.

For immunostaining of target proteins, cells were rinsed with PBS, fixed in 4% paraformaldehyde (PFA) for 20 minutes at room temperature, and permeabilized in 0.1% Triton X-100 in PBS for 1 hour. Cells were then blocked with 1% bovine serum albumin (BSA) in PBS for 1 hour. After three washes with PBS (5 minutes each on a rocker), coverslips were incubated overnight at 4°C with primary antibodies diluted in blocking buffer. The following primary antibodies were used: anti-ZO-1 (33-9100, Thermo Fisher Scientific), anti-ICAM1/CD54 (ab171123, Abcam), recombinant anti-VCAM1 (ab134047, Abcam), anti-VE-Cadherin (ab282277, Abcam), anti-Integrin αV (ab179475, Abcam), anti-p21 (ab188224, Abcam), anti-p27 (ab32034, Abcam), anti-p53 (648501, BioLegend), and Lamin B1 (sc-374015, Santa Cruz Biotechnology).

After primary antibody incubation, cells were washed three times in PBS and incubated with the appropriate secondary antibodies for 1 hour at room temperature: Alexa Fluor® 488 or 594 AffiniPure Goat Anti-Mouse IgG (115-545-146/115-585-146, Jackson ImmunoResearch) and Alexa Fluor® 488 or 594 AffiniPure Donkey Anti-Rabbit IgG (711-545-152/711-585-152, Jackson ImmunoResearch). Following secondary antibody incubation, nuclei were counterstained with DAPI (D9542, Sigma-Aldrich; 1:2000 in Milli-Q water) for 10 minutes at room temperature, then coverslips were washed in Milli-Q water.

Coverslips were mounted on Superfrost Plus microscope slides using Fluoroshield™ mounting medium (F6182, Sigma-Aldrich) and stored at 4°C. Imaging was performed using a LionHeart FX microscope (BioTek) at 10X, 20X, and 40X objectives. Images from 10X and 20X magnification were used for quantitative analysis, while 40X images were used for illustrative purposes. For comparative analysis, identical microscope settings (exposure time, gain, and LED illumination) were applied to paired control and test samples within each individual experiment.

### Western Blotting

First, bEnd.3 cells were cultured in 10-cm culture dishes and grown in DMEM medium. Upon reaching approximately 70% confluence, cells were treated with isoDGR-peptide or isoD-peptide (control) at 50 μg/mL, prepared in autoclaved 1X PBS) for 24-48 hours. For positive controls, a subset of cells was primed with lipopolysaccharide (LPS, 1 μg/mL, L2630, Sigma-Aldrich) for 4-6 hours. Untreated cells served as negative controls.

After treatment, cells were washed with cold PBS and lysed on ice in RIPA buffer (150 mM NaCl, 1% Triton X-100, 0.5% sodium deoxycholate, 0.1% SDS, 50 mM Tris-HCl, pH 8.0) supplemented with 0.1% protease inhibitor cocktail (ThermoFisher Scientific). Lysates were collected, sonicated briefly, and centrifuged at 12,000g for 15 minutes at 4°C. Protein concentrations were determined using the Bradford assay (Bio-Rad). Equal amounts of protein (30-40 μg per lane) were separated on 10-20% SDS-polyacrylamide gels and transferred to PVDF membranes (Immobilon-P, Millipore).

Membranes were blocked for 1 hour at room temperature with 5% BSA or 5% skim milk in TBST (0.1% Tween-20 in Tris-buffered saline), and then incubated overnight at 4°C with primary antibodies diluted 1:1000 in blocking buffer. The following antibodies were used: anti-p27 (ab32034, Abcam), anti-p21 (ab188224, Abcam), Lamin B1 (sc-374015, Santa Cruz Biotechnology), anti-ZO-1 (33-9100, Thermo Fisher Scientific), anti-ICAM1 (ab171123, Abcam), anti-VCAM1 (ab134047, Abcam), anti-VE-Cadherin (ab282277, Abcam), anti-Integrin αV (ab179475, Abcam), anti-β-actin (4970L, Cell Signaling Technology), and anti-GAPDH (ab8245, Abcam).

After three 5-minute washes in TBST, membranes were incubated for 1 hour at room temperature with horseradish peroxidase (HRP)-conjugated secondary antibodies (HRP Goat anti-Mouse IgG, IgM, IgA (A-10668, ThermoFisher Scientific) or HRP Goat anti-Rabbit IgG (AS014, ABclonal), diluted in blocking buffer as recommended by the manufacturer). Detection was performed using Pierce™ ECL Western Blotting Substrate (Thermo Fisher Scientific), and chemiluminescent signals were captured using the Chemidoc Imaging System (Bio-Rad).

Band intensities were quantified using Fiji/ImageJ software (NIH). Signals were normalized to housekeeping proteins (β-actin or GAPDH) within each lane. Relative protein expression was calculated as fold change compared to the negative control condition.

### Quantitative Real-Time PCR (qRT-PCR)

Total RNA was extracted from bEnd.3 cells using the RNeasy Mini Kit (Qiagen) with on-column DNase I treatment. RNA concentration and purity were measured by NanoDrop. First-strand cDNA was synthesized from 1 μg RNA using the SuperScript™ IV First-Strand Synthesis System (Invitrogen). qRT-PCR was performed using PowerTrack™ SYBR Green Master Mix (Thermo Fisher Scientific) on a QuantStudio™ 3 Real-Time PCR System. Each 20 μL reaction contained 10 μL master mix, 0.5 μM each primer, and 1 μL cDNA. The cycling protocol was 95°C for 2 min, followed by 40 cycles of 95°C for 15 s and 60°C for 1 min. Melt curve analysis confirmed specificity. Gene expression was quantified from standard curves and normalized to GAPDH mRNA. Results represent five biological replicates, analyzed in duplicate. The primers used are listed in Table S1.

### CellROX and MitoSOX Assays

To assess intracellular and mitochondrial reactive oxygen species (ROS), bEnd.3 cells were seeded onto sterile glass coverslips in 6-well plates and cultured overnight in DMEM cell culture medium at 37°C and 5% CO₂, allowing cells to reach approximately 70% confluence. The next day, culture medium was replaced, and cells were treated with isoDGR-peptide at concentrations of 50 μg/mL or 100 μg/mL, or with 0.2 μM doxorubicin (44583, Sigma-Aldrich) as a positive control; untreated cells served as negative controls. Treatments were maintained for 96 hours, with medium refreshed after 48 hours. At the end of the treatment period, ROS probes were applied. For total ROS detection, CellROX Green reagent (Thermo Fisher Scientific) was freshly diluted in DMSO to a final concentration of 0.5 μM and added directly to the wells. For mitochondrial ROS, MitoSOX Green (M36005, Thermo Fisher Scientific) was similarly prepared and applied at 0.5 μM. Cells were incubated for 30 minutes at 37°C in the dark. Following incubation, wells were gently washed with warm PBS and cells were fixed with 4% paraformaldehyde for 15 minutes at room temperature. A blank well containing cells but no dye was included in each experiment to control for autofluorescence. Coverslips were mounted using Fluoroshield mounting medium (F6182, Sigma-Aldrich). Fluorescence images were acquired using a LionHeart FX microscope (BioTek) at 10X or 20X magnification for quantitative analysis and at 40X for representative structural imaging. ROS signal intensity was quantified using ImageJ software, and all quantifications were performed by investigators blinded to treatment groups.

### ROS Quantification Assay

Dynamic changes in intracellular ROS were quantified using the dichlorofluorescin diacetate (DCFH-DA) assay as previously described (Ávila et al., 2024)^44^. Briefly, bEnd.3 cells were cultured and treated as described above. Following treatment, cells were harvested by trypsinization, washed with PBS, and homogenized in 20 mM sodium phosphate buffer (pH 7.4) containing 140 mM KCl. Lysates were centrifuged at 3,500 × g for 10 minutes at 4°C, and the supernatant was collected for analysis. Immediately prior to measurement, 2’,7’-dichlorofluorescin diacetate (DCFH-DA; D6883, Sigma-Aldrich) was freshly prepared in DMSO and added to the supernatant at a final concentration of 20 μM. Samples were incubated at 37°C, and fluorescence was measured at excitation/emission wavelengths of 480/520 nm using a DeNovix fluorometer at multiple time points. ROS levels were normalized to total protein content as determined by the Bradford assay. All conditions were performed in triplicate, and investigators were blinded to group assignments during quantification.

### Colony Formation Assay

Colony formation was assessed using a standard crystal violet staining protocol. bEnd.3 cells were seeded at a density of 200 cells per well into 6-well tissue culture plates (Corning) in DMEM medium. Cells were allowed to adhere overnight at 37°C in a humidified 5% CO₂ incubator. The following day, wells were treated with either isoDGR-peptide dissolved in sterile 1X PBS at a final concentration of 50 μg/mL or 100 μg/mL, or left untreated (control).

After treatment, cells were cultured for an additional 7 days, with medium changed every 48-72 hours. At the end of the incubation, wells were washed with 1X PBS and colonies were fixed using 4% paraformaldehyde (PFA; Sigma-Aldrich) for 20 minutes at room temperature. Following fixation, cells were washed three times with PBS and stained with 1 mL of 0.5% crystal violet solution (C0775, Sigma-Aldrich) in 25% methanol for 10-20 minutes. Plates were then rinsed thoroughly with Milli-Q water to remove excess dye and air-dried.

Colonies were imaged at 4X magnification using a brightfield microscope (LionHeart FX, BioTek). Colony number and area were quantified in at least three independent experiments per condition. Only colonies consisting of ≥50 cells were counted and included in the analysis, in accordance with standard protocols. Colony numbers were quantified using ImageJ software.

### MTT Cell Viability Assay

Cell viability in response to isoDGR-peptide exposure was assessed using the MTT assay. bEnd.3 cells were seeded in 96-well plates (Corning) and cultured in DMEM medium as described above. Cells were grown to 70% confluence prior to treatment. IsoDGR-peptide was added to culture media at final concentrations of 1, 5, 10, 25, 50, or 100 μg/mL. Each condition was performed in triplicate, and wells without cells (media only) served as the blank control. Cells were incubated with treatments for 24-96 hours, depending on the experimental design. At the endpoint, cell viability was assessed by adding MTT reagent (Thiazolyl Blue Tetrazolium Bromide; 475989, Sigma-Aldrich) at a final concentration of 0.5 mg/mL to each well. Plates were returned to the incubator for 30 min to 2 hr to allow formation of purple formazan crystals, which reflect metabolically active cells. Following incubation, culture media was gently removed without disturbing the crystals, and 150 mL of DMSO was added per well to fully solubilize the formazan product. Absorbance was measured at 570 nm (reference 630 nm) using a Cytation 5 plate reader or in a DeNovix spectrophotometer. All readings were normalized by subtracting the mean absorbance of the blank wells. Relative cell viability was expressed as a percentage of untreated controls. All experiments were conducted in at least three independent replicates. Investigators were blinded to treatment conditions during absorbance quantification.

### Senescence-Associated β-Galactosidase (SA-β-gal) Staining

Senescence was induced in mouse brain microvascular endothelial bEnd.3 cells by treatment with isoDGR-peptide [GC(isoD)GRCGK] (50 μg/mL) or doxorubicin (44583, Sigma-Aldrich; 0.1-100 μM) as a positive control. Cells were plated onto plain glass coverslips in DMEM medium and maintained at 37°C in a humidified atmosphere with 5% CO₂. Treatments were applied once cells reached approximately 70% confluence (typically day 2 post-plating) and continued for 48-96 hours. For detection of senescence-associated β-galactosidase (SA-β-gal) activity, cells were washed with PBS and fixed in 4% paraformaldehyde (PFA) in PBS for 3-5 minutes. After three washes in PBS, cells were incubated with X-Gal staining solution (0.1% X-Gal, 5 mM potassium ferricyanide, 5 mM potassium ferrocyanide, 2 mM MgCl₂, 150 mM NaCl, and 40 mM sodium phosphate/citric acid buffer, pH 6.0) for 9-20 hours at 37°C in a non-CO₂ incubator. Blue staining was monitored periodically by brightfield microscopy. Once optimal staining was observed, the reaction was terminated by washing cells three times in Milli-Q water for 5 minutes each. Coverslips were immediately imaged.

### Statistical Analysis

All statistical analyses were performed using GraphPad Prism version 9.3 (GraphPad Software, San Diego, CA). Data were assessed for normality using the D’Agostino-Pearson omnibus or Shapiro-Wilk tests, as appropriate. For comparisons between two groups with normally distributed data, unpaired two-tailed Student’s t-tests were applied. For analysis of more than two groups, one-way analysis of variance (ANOVA) followed by Tukey’s post hoc multiple comparisons test was used. If data were not normally distributed, the Kruskal-Wallis test with Dunn’s multiple comparisons post hoc test was applied. Repeated measures data were analyzed using repeated measures ANOVA. Quantitative results are presented as mean ± standard error of the mean (SEM), unless otherwise indicated. P values less than 0.05 were considered statistically significant. All statistical tests were two-sided.

## DATA AVAILABILITY

The RNA-seq data supporting this study have been deposited in NCBI GEO under accession number GSE301193, where they are accessible for download. All remaining data are included within the main manuscript or its supplementary materials.

## ACKNOWLEDGMENTS

This work was supported in part by the Canadian Institutes of Health Research Tier1 Canada Research Chair (CRC-2020-00263), Canadian Institutes of Health Research Project Grant (PJT-186091), The Natural Sciences and Engineering Research Council of Canada Discovery Grant (RGPIN-2023-04304), Canada Foundation for Innovation Grant (41454 and 44115), Ontario Research Fund, start-up research grant from Brock University, and Singapore National Medical Research Council (NMRC/OFIRG/0003/2016).

## AUTHOR CONTRIBUTIONS

R.P., K.P., and S.K.S. conceived and designed the project; R.P., K.P., S.C.N., R.I., J.N., B.D., M.K., Y.T., and M.H.L., performed the experiments and data analysis; R.P. and S.K.S. wrote the manuscript; B.E.F., R.E.K.M., E.L.T., P.K., D.K., R.N.K. and N.E.M. provided resources/equipment; R.E.K.M., E.L.T., P.K., N.E.M. and S.K.S contributed grants/reagents/materials/analysis tools; S.K.S supervised the project. All authors revised the manuscript.

---

**Table S1.**
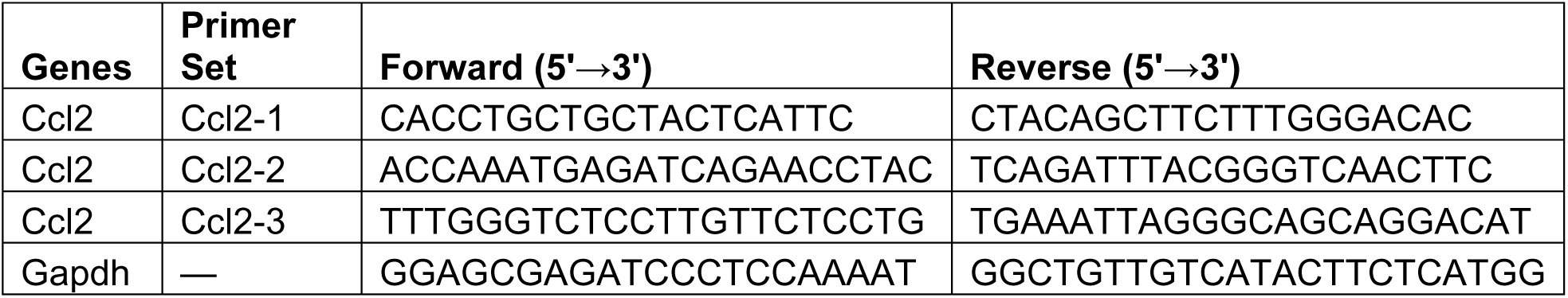
Primer Sequences for qRT-PCR Amplification of Mouse Ccl2 and Gapdh Genes.

**Figure S1.**
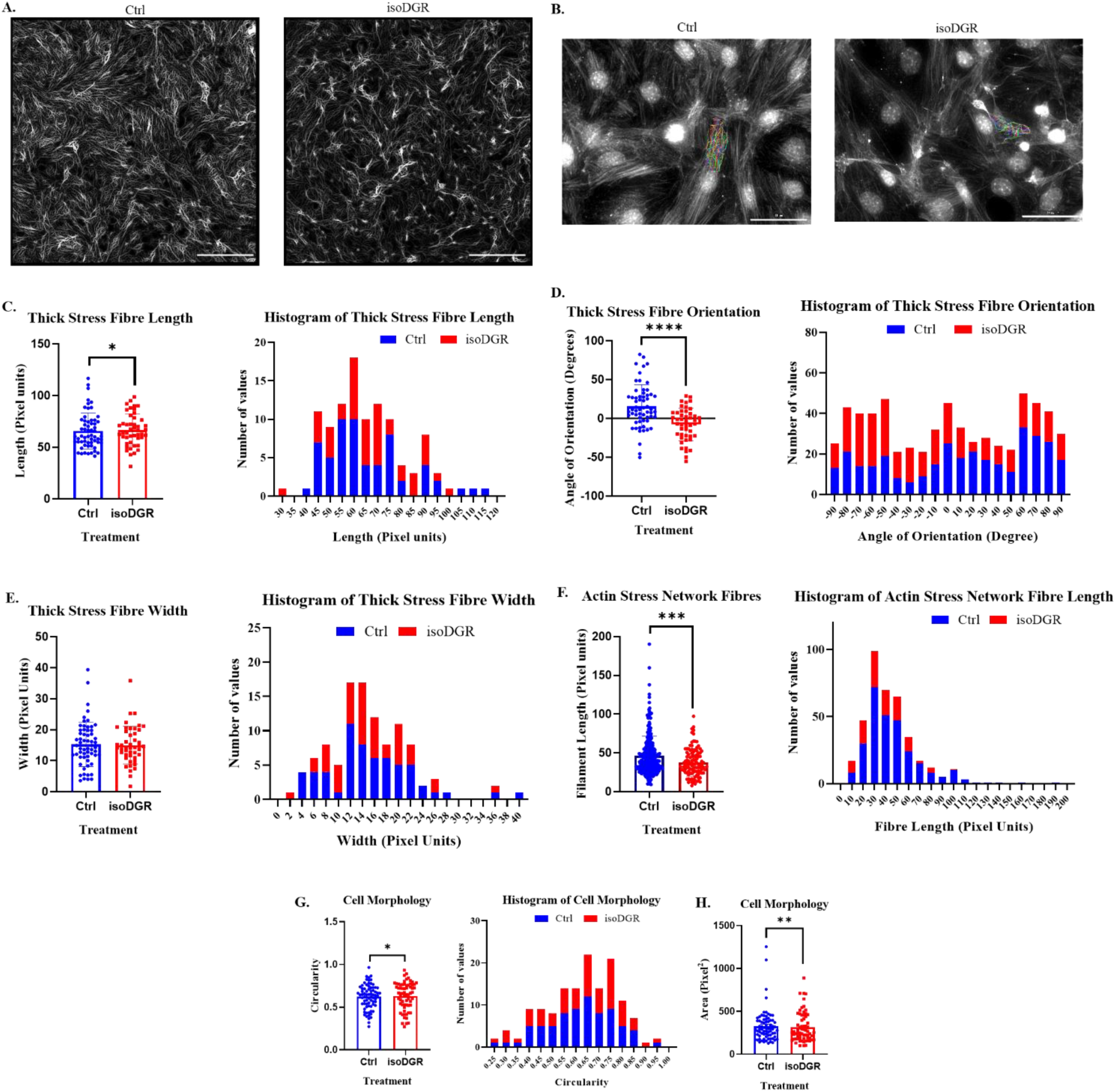
Automated detection, tracing, and quantitative analysis of actin stress fibers and cell morphology in bEnd.3 cells under control and isoDGR-treated conditions using the SFEX method^43^. (A) Actin stress fiber network detection: Representative fluorescence image of F-actin cytoskeletal organization following processing with the Stress Fiber Extractor (SFEX) algorithm. Scale bar = 100 μm. (B) Thick stress fiber tracing: High-intensity actin bundles were automatically segmented and reconstructed using SFEX. Scale bar = 30 μm. (C) Total fiber length: Scatter plot and histogram showing total actin fiber length per cell in control (blue) and isoDGR-treated (red) groups. (D) Fiber orientation: Scatter plot and histogram comparing average fiber orientation per cell between groups. (E) Fiber width: Scatter plot and histogram comparing mean actin fiber width per cell between groups. (F) Network fiber length: Scatter plot and histogram comparing total F-actin network fiber length per cell between groups. (G) Cell circularity: Scatter plot and histogram comparing cell circularity between groups. (H) Cell area: Scatter plot comparing cell area (in pixels²) between groups. Data in panels C-H are presented as individual values (scatter plots) and distribution profiles (histograms) for each group. Statistical comparisons were performed between control and isoDGR-treated cells; p < 0.05 considered significant (*p < 0.05; **p < 0.01; ***p < 0.005; ****p < 0.001).

